# 4D bioprinted self-folding scaffolds enhance cartilage formation in the engineering of trachea

**DOI:** 10.1101/2023.12.06.570378

**Authors:** Irene Chiesa, Alessio Esposito, Giovanni Vozzi, Riccardo Gottardi, Carmelo De Maria

## Abstract

Trachea defects that required surgical interventions are increasing in number in the recent years, especially for pediatric patients. However, current gold standards, such as biological grafts and synthetic prothesis, do not represent an effective solution, due to the lack of mimicry and regeneration capability. Bioprinting is a cutting-edge approach for the fabrication of biomimetic scaffold to empower tissue engineering toward trachea replacement. In this study, we developed a self-folding gelatin-based bilayer scaffold for trachea engineering, exploiting the 4D bioprinting approach, namely the fabrication of dynamic scaffolds, able to shape morph in a predefined way after the application of an environmental stimulus. Indeed, starting form a 2D flat position, upon hydration, this scaffold forms a closed tubular structure. An analytical model, based on Timoshenko’s beam thermostats, was developed, and validated to predict the radius of curvature of the scaffold according to the material properties and the scaffold geometry. The 4D bioprinted structure was tested with airway fibroblast, lung endothelial cells and ear chondral progenitor cells (eCPCs) toward the development of a tissue engineered trachea. Cells were seeded on the scaffold in its initial flat position, maintained their position after the scaffold actuation and proliferated over or inside it. The ability of eCPCs to differentiate towards mature cartialge was evaluated. Interestingly, real-time PCR revealed that differentiating eCPCs on the 4D bioprinted scaffold promote healthy cartilage formation, if compared with eCPCs cultured on 2D static scaffold. Thus, eCPCs can perceive scaffold folding and its final curvature and to react to it, towards the formation of mature cartilage for the airway.

## 1. Introduction

The trachea is a tube-shaped organ that connects the larynx to the lungs. It allows for respiration, cleaning and humidifying the inspired air, and plays a key role in speaking [1], [2]. This organ possesses a hierarchical graded structure consisting, from the lumen outward, of epithelium, basement membrane, connective tissue, smooth muscle, and hyaline cartilage, which is organized in 18 to 22 C-shaped open rings. Trachea defects (*e.g.,* congenital defects, stenosis, tumours, trauma) that are complex in shape or involved more than 50% of its length (30% in paediatric patients [3]) require the surgical removal of the trachea and its replacement with auto- or allograft or with synthetic implants [4], [5]. However, the use of autograft is limited by the lack in the human body of sufficient appropriate replacement tissue to restore the authentic 3D composite structure of the trachea, whereas allografts are limited by the host immune response [6]–[9]. On the other side, the use of synthetic prosthesis, typically made of silicone, rarely leads to tissue regeneration and are often short-lived for mismatch with the surrounding native tissue [10]. In this context, scaffold-based tissue engineering offers a cutting-edge method for the engineering of human tissues, representing a promising approach for trachea reconstruction. However, conventional fabrication techniques such as salt leaching and solvent casting lack the fine control over scaffold geometry and material spatial distribution necessary for tracheal engineering. Hence, researchers turned to bioprinting, namely the use of additive manufacturing (AM) technologies to fabricate 3D constructs with high spatial control and designed to interact with physiological systems at cellular levels [11], [12].

For instance, Ke *et al*. [13] used bioprinting to manufacture a tracheal scaffold based on polycaprolactone (PCL) combined with a mesenchymal stem cells loaded hydrogel to form a biphasic construct of cartilage and smooth muscle. Similarly, Park *et al.* [14] exploited a dual-extruder bioprinter equipped with a rotating spindle to fabricate a multilayered cylindrical trachea scaffold with a PCL backbone and two separate alginate-based confined cellular compartments for chondrocytes and epithelial cells.

However, although highly promising, the overmentioned 3D bioprinted scaffolds are not able to closely mimic the native tissue dynamics of native tissue, including the trachea, due to their static behaviours.

In this scenario, the recent years, four dimensional (4D) bioprinting has been developed, in which the bioprinted constructs are no longer static objects but programmable active structures that are able to shape-transform remotely upon the application of an external stimulus [15], [16]. In 4D bioprinting, passive and active (also referred to as smart or responsive) materials are deliberately arranged in space, so that chemical/physical variation at the materials level will result in a macroscopic shape transformation of the construct [17]–[19].

Compared to static constructs, 4D bioprinted ones offer the exciting ability to mimic tissue dynamics, to provide stimuli promoting cell activity and differentiation, to adapt to the 3D complex shape of human body, and to reshape over time for minimally invasive applications and [20]–[22]. Moreover, scaffold fabrication and cell seeding can be simplified by 4D bioprinting which can be fabricated in their initial 2D position and then reach their final 3D conformation after cell incorporation and actuation.

For example, Kim *et al*. [23] exploited 4D bioprinting to fabricate a silk-based self-folding scaffold for partial tracheal replacement. The construct, comprising both chondrocytes and turbinate-derived mesenchymal stem cells, was implanted in a rabbit model with a damaged trachea for 8 weeks, and integrated well with the formation of novel epithelium and the establishment of cartilage layers in the predicted sites.

In this work, we exploited the 4D bioprinting approach to fabricate a self-folding bilayer scaffold for trachea engineering. The self-folding over-time was achieved thanks to the differential swelling properties that the two layers of the scaffold possess, resulting in the formation of a close tubular structure. To predict the radius of curvature of the scaffold from materials parameters and scaffold geometry, we developed an analytical model based on Timoshenko’s bimetal thermostats. Different cell types were easily seeded on either side of the scaffold in its initial flat 2D configuration that, after hydration, self-folded into a cell-laden tubular structure with distinct cellular compartments (*i.e*., cartilage rings, fibroblast, epithelium layers), providing the cells with a proper environment for their proliferation and differentiate. Finally, we studied the formation of a mature cartilage ring derived from the differentiation of human ear cartilage progenitor cells on the 4D bioprinted scaffolds.

## 2. Materials and Methods

### 2.1 Material preparation and characterization

Type A gelatin from porcine skin (Sigma-Aldrich), crosslinked by (3-Glycidoxypropyl) trimethoxysilane (GPTMS, Sigma-Aldrich) was selected as bulk material for this study. The high-swelling layer of the scaffold was made with a 15% w/v gelatin in phosphate-buffered saline solution (PBS, Sigma-Aldrich) 1X crosslinked with 92 μl/g of GPTMS (GPTMS-GEL-15). The low-swelling layer was made with a 5% w/v gelatin solution in PBS 1X crosslinked with 368 μl/g of GPTMS (GPTMS-GEL-5) [24]. Each solution was prepared by dissolving the gelatin in PBS 1X for 90 min at 60°C on a stirring plate, and then adding the GPTMS keeping stirring for 40 min at 60°C. Monolayer films of either material were fabricated by solvent and dried for 24 h at room temperature and characterized in terms of swelling coefficient and elastic modulus.

Swelling tests were performed on rectangular films (n = 10). Briefly, dimensions of each sample were measured at dry state. The x-y dimensions were acquired via image analysis through ImageJ, while thickness (dimension along z) was acquired with a micrometer. Then, films were placed in a 24-well plate, submerged with 1 ml PBS 1X and placed at 37°C. Dimensions of each sample were evaluated after 24h, and the swelling coefficient in each dimensionwas evaluated as in Eq. 1.

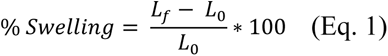

where L_f_ is the dimension after 24h, and L_0_ is the dimension at dry state.

Elastic modulus of each material, in both dried and hydrated state (*i.e*., 24 h in PBS 1X), was evaluated (n = 3) by uniaxial tensile testing using a Zwich-Roell z005 machine equipped with a 100 N load cell at a strain rate of 10% min^-1^. The elastic modulus was calculated as the slope of the linear region in the stress-strain curve obtained from the tensile test.

### 2.2 Fabrication process of the scaffolds via 4D bioprinting

3.5 ml of GPTMS-GEL-15 solution was poured into Petri Dishes (Ø = 60 mm) that were carefully sealed with Parafilm and kept at 37°C for 36 h, thus allowing the crosslinking process to begin without the complete evaporation of the solvent. Then, 3-layered lines of GPTMS-GEL-5 were bioprinted on the GPTMS-GEL-15 film via piston-driven extrusion-based bioprinting exploiting a custom-made bioprinter, previously described in [12], [25], with the following printing parameters: print speed = 5 mm/s, needle diameter = 0.4 mm, layer height = 0.1 mm, volumetric flow = 0.4 mm^3^/s. Then, the bilayer scaffolds were dried for 48 hours at room temperature and finally cored into 10 mm x 5 mm rectangular samples. The fabrication technique is summarized in Figure 1. Film thickness, line distances, and line width were evaluated via image analysis with ImageJ.

**Figure 1:**
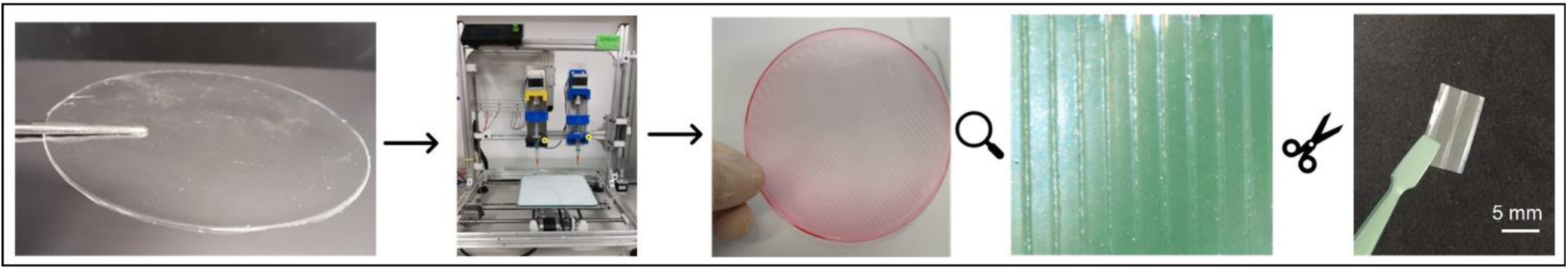
Main fabrication steps of the self-folding scaffolds: firstly, the high-swelling layer was fabricated by solvent casting. Then, 3-layered lines of the low-swelling material were 4D bioprinted via extrusion-based bioprinting on the previous layer. After drying, 10 mm x 5 mm samples were cored from the obtained films. Food dyes were added in the material only visualization purpose.

### 2.3 Scaffold characterization

The scaffold capability to self-fold into a tube in aqueous solution was verified by dipping the dried scaffold in PBS 1X. The ability to control the folding direction according to the bioprinted line orientation was verified (Figure 2A). Then, for each configuration (*i.e.*, bioprinted lines parallel to the long side and bioprinted lines parallel to the short side) the final length, internal, and external diameter of the actuated tube were evaluated via image analysis with ImageJ (n = 5). The folded scaffolds thickness was evaluated as the difference between the external and internal diameter of each scaffold.

**Figure 2:**
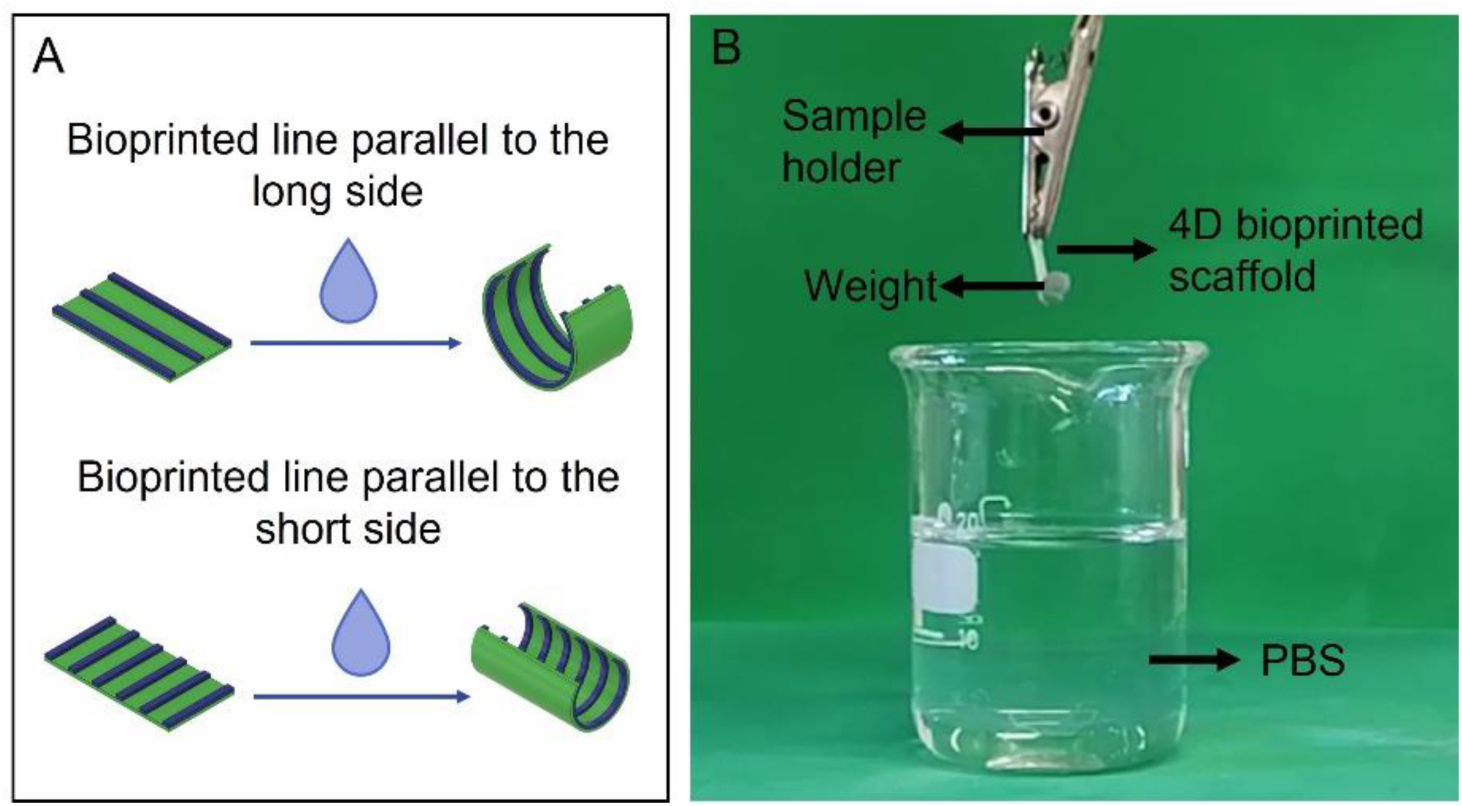
Scaffold characterization. A) The ability of the 4D bioprinted scaffold to self fold according to the bioprinted line orientation was verified dipping the scaffolds in PBS 1X. B) Experimental set up for the evaluation of the force exerted by the scaffold during its self-folding.

The force exerted by the scaffold during its shape-shifting was also measured. Briefly, hollow cylinders with predefined weight (*i.e*., 20 to 80 mg with a step of 10 mg) were designed via Fusion360 and fabricated via stereolithography (Form2 by Formlabs, using the Draft resin). Then, the cylinder was sutured on one edge of the samples, whereas the other edge was clamped to a holder (Figure 2B). The samples were then dipped in PBS 1X and allowed to self-fold. The maximum weight that allows the tube to self-fold were used to measure the exerting force exploiting the Archimede’s principles, as shown in Eq. 2.

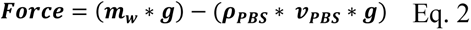

where *m_w_* is the mass of the cylindrical weight, *g* is the gravity acceleration, *ρ_PBS_* is the density of the PBS 1X and *v*_PBS_ is the volume of the displaced fluid, that is equal to the volume of the cylinder.

### 2.4 Analytical model to predict the self-folding of the scaffold

#### 2.4.1 Model hypothesis, derivation, and validation

Starting from Timoshenko’s model [26], that deals with bimetal beams under thermal expansion, an analytical model to predict the curvature of the self-folding scaffold was developed. The geometry under analysis can be schematized as a large parallelepiped (solid layer, GEL-GPTMS-15) with a series of N equally spaced smaller parallelepipeds on its top surface (bioprinted layer, GEL-GPTMS-5), as shown in Figure 3A. The model aims to solve the so-called forward problem, *i.e.,* to relate the final radius of curvature of the structure (ρ) to: i) the initial geometry of the scaffold; ii) the properties of the involved materials; iii) the applied stimulus (Figure 3B).

**Figure 3:**
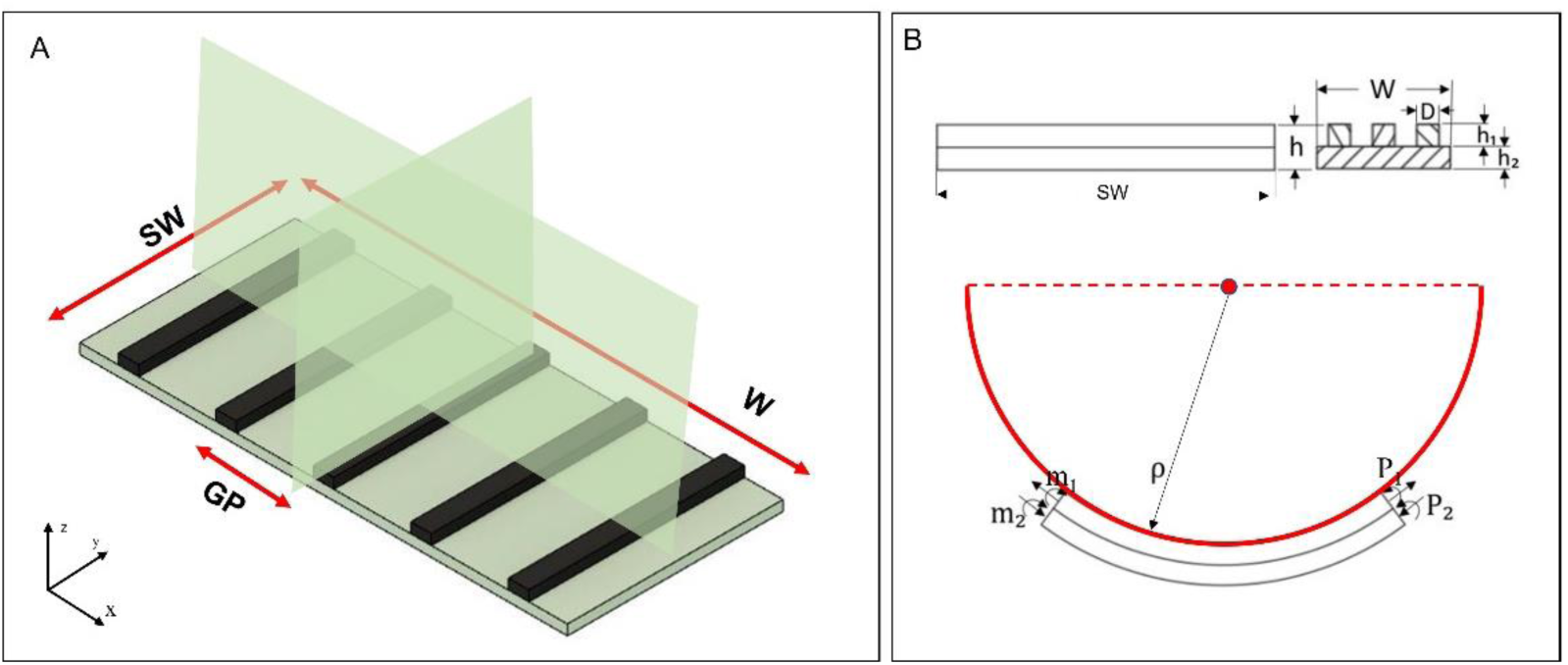
A) Schematic of the modelled bilayer structure, where the planes of symmetry are reported (green). B) Geometrical value of interest in the model, with the involved forces (P_1_, P_2_) and torques (m_1_, m_2_).

The input parameters for the model are reported and described in Table 1. Due to the periodicity of the modeled structure, a duty cycle (*i.e*., the ratio between the width of the bioprinted line and the geometrical period (GP) defined as the sum of the width of a bioprinted line and the width of the interlaying space) can be defined as in Eq. 3.

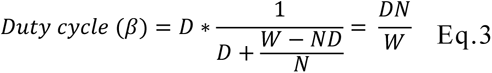

**Table 1:**
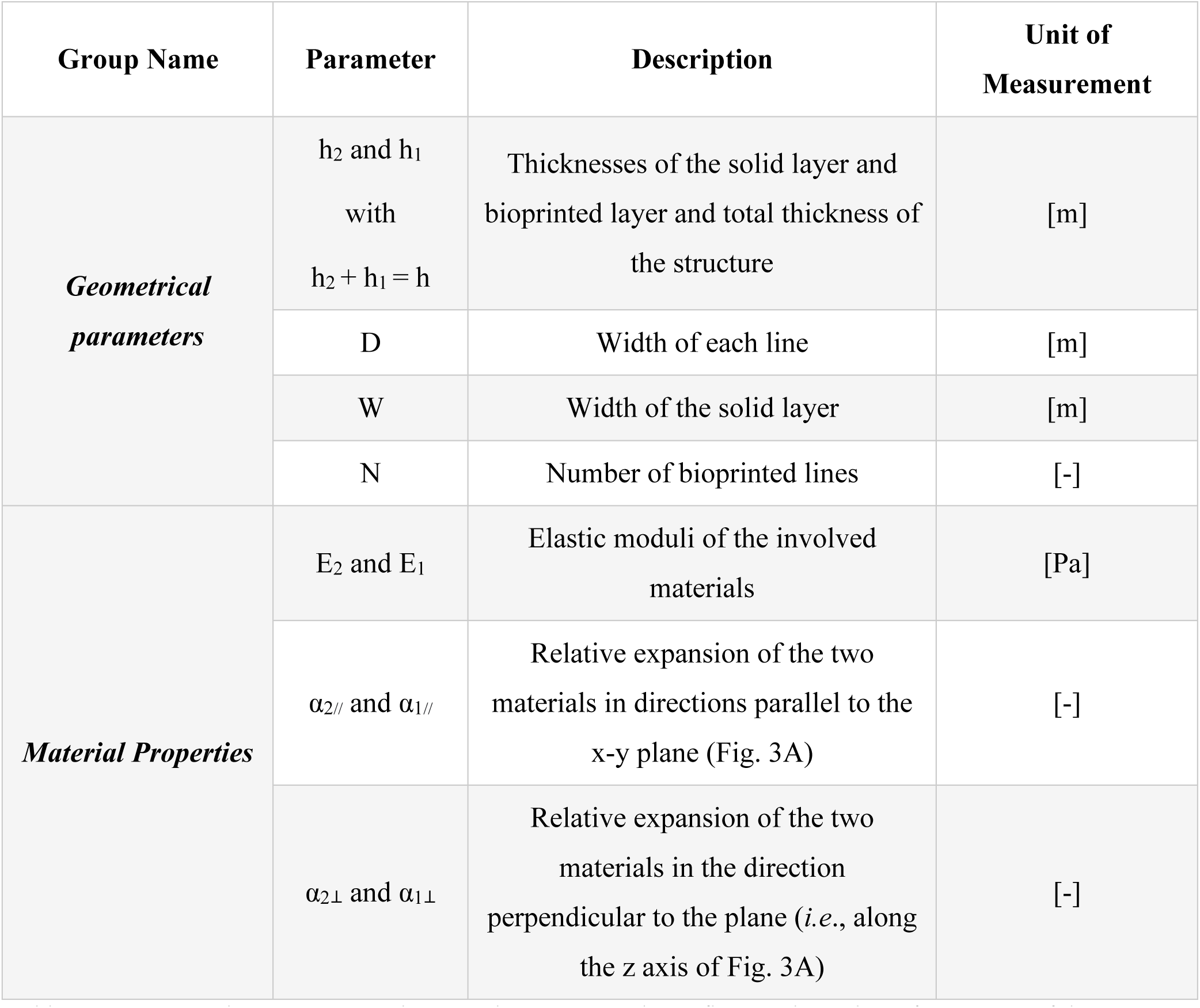
Geometrical parameters and materials properties that influence the radius of curvature of the structure.

| Group Name | Parameter | Description | Unit of Measurement |
| --- | --- | --- | --- |
| **Geometrical parameters** | $h_2$ and $h_1$<br>with<br>$h_2 + h_1 = h$ | Thicknesses of the solid layer and bioprinted layer and total thickness of the structure | [m] |
|  | D | Width of each line | [m] |
|  | W | Width of the solid layer | [m] |
|  | N | Number of bioprinted lines | [-] |
| **Material Properties** | $E_2$ and $E_1$ | Elastic moduli of the involved materials | [Pa] |
| | $\alpha_{2//}$ and $\alpha_{1//}$ | Relative expansion of the two materials in directions parallel to the x-y plane (Fig. 3A) | [-] |
| | $\alpha_{2\perp}$ and $\alpha_{1\perp}$ | Relative expansion of the two materials in the direction perpendicular to the plane (*i.e.*, along the z axis of Fig. 3A) | [-] |

The model hypothesis and derivation are described in detail in the supplementary information (SI) 1 and involve the equilibrium equations of force and torque, the constituent equations, and the continuity equations.

The model was validated through a parametric sensitivity analysis, focused on three parameters: i) the total thickness of the scaffold (h); ii) the duty cycle (β), which can be changed by varying the bioprinted pattern; and iii) the total dimension of the scaffold, achieved by the variation of W, L, h. The validation of the model is described in SI2.

### 2.5 Biocompatibility of the 4D bioprinted scaffold

#### 2.5.1 Cell culture

Three different cells sources were separately employed. Immortalized human airway fibroblasts (AFs) were kindly provided by Dr. Susan L. Thibeault from University of Wisconsin-Madison. AFs will be always seeded on the internal-to-be layer of the scaffold, and, when not otherwise stated, they were cultured in Dulbecco’s Modifies Eagle Medium (DMEM) supplemented with 10% v/v fetal bovine serum (FBS), 2% v/v penicillin/streptomycin/fungizone (PSF) and 2% v/v of nonessential amino acids.

Immortalized non-tumorigenic human lung epithelial cells (ECs), were purchased from ATCC. ECs will be always seeded on the internal-to-be layer of the scaffold, and, when not otherwise stated, they were cultured in Ham’s F12 medium supplemented with 4% v/v FBS, 2% v/v PSF, 1.5 g/L sodium bicarbonate, 2.7 g/L glucose, 2.0 mM L-glutamine, 0.1 mM nonessential amino acids, 0.005 mg/ml insulin, 0.001 mg/ml transferrin and 500 ng/ml hydrocortisone.

Human ear chondral progenitor cells (eCPCs) were extracted by Dr. Soheila Ali Akbari Ghavimi from patients’ samples provided by the Children’s Hospital of Philadelphia (Children’s Hospital of Philadelphia IRB 20-017817, 20-017551, 20-017287). eCPCs will be always seeded on the external-to-be layer of the scaffold, and, when not otherwise stated, they were cultured in growth medium made of DMEM:F12 medium supplemented with 10% FBS, 2% PSF, 1 mg/ml L/glucose, 2.34 mg/ml (4-(2-hydroxyethyl)-1-piperazineethanesulfonic acid), 0.5 μl/ml Ascorbic Acid, 0.5 μl/ml fibroblast growth factor 2 and 0.1 μl/ml transforming growth factor beta 1 (TGFβ1).

#### 2.5.2 Scaffold seeding and culture

4D bioprinted scaffolds in their flat initial position were sterilized by 30 mins of ultraviolet (UV) light on each side. Then, they were place in a 12 well plate under the central cavity of a custom holder (SI 4). 20k cells suspended in 100 μl of cell medium were delivered inside the holder cavity. After 90 mins, 1 ml of medium was added outside the holder and an additional 100 μl inside the holder cavity. The three cell populations mentioned above were separately seeded: i) AFs were seeded on the internal-to-be-layer, ii) ECs were seeded on the internal-to-be-layer; iii) eCPCs were seeded on the external to be layer. Scaffolds were then incubated at 37°C and 5% CO_2_ overnight to allow cells to adhere to the scaffold. The next day, the holders were removed, allowing the scaffolds to self-fold, and 2 ml of fresh medium were added. Scaffolds were then cultured for 10 days, renewing the medium every 2 days. Alamar Blue (Invitrogen) assays was performed on the scaffolds (n = 3) at 24 h, 72 h, 7 days, and 10 days after the seeding. Briefly, 10% v/v of Alamar Blue solution in PBS 1X was added to the samples (3 ml for sample) and incubate for 2h in the dark. Then, fluorescence was read (ex/em ∼ 560/590 nm) with a plate reader (Synergy HT). A standard curve for the estimation of cells number was created using different number of cells (*i.e.*, 10^3^, 5*10^3^, 10^4^, 5*10^4^, 10*10^5^, 2*10^5^ cells) cultured on tissue culture plates. Seeding efficacy after the scaffold actuation was estimated for each cell type and scaffold orientation as reported in Eq. 4.

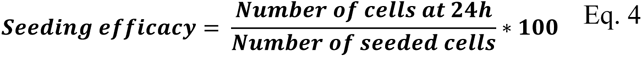

### 2.6 Development of tracheal cartilage rings

eCPCs were seeded on the scaffold and the cell seeded scaffolds were actuated 24 h after seeding, as described in 2.5.2. Then, constructs were cultured for 7 days in growth medium to allow cells to proliferate and cover the entire scaffolds. At this point, the constructs were switched to chondrogenic medium (DMEM supplemented with 2% PSF, 10ng/ml TGF-β3, 50 μM ascorbic acid, 10 nM dexamethasone and 23μM proline) and cultured for 21 additional days. A schematic of the experiment set up is shown in Figure 4A. 2D passive scaffolds fabricated via solvent casting with GEL-GPTMS-15 were used as control, sterilized, and seeded with the same protocols Figure 4B.

**Figure 4:**
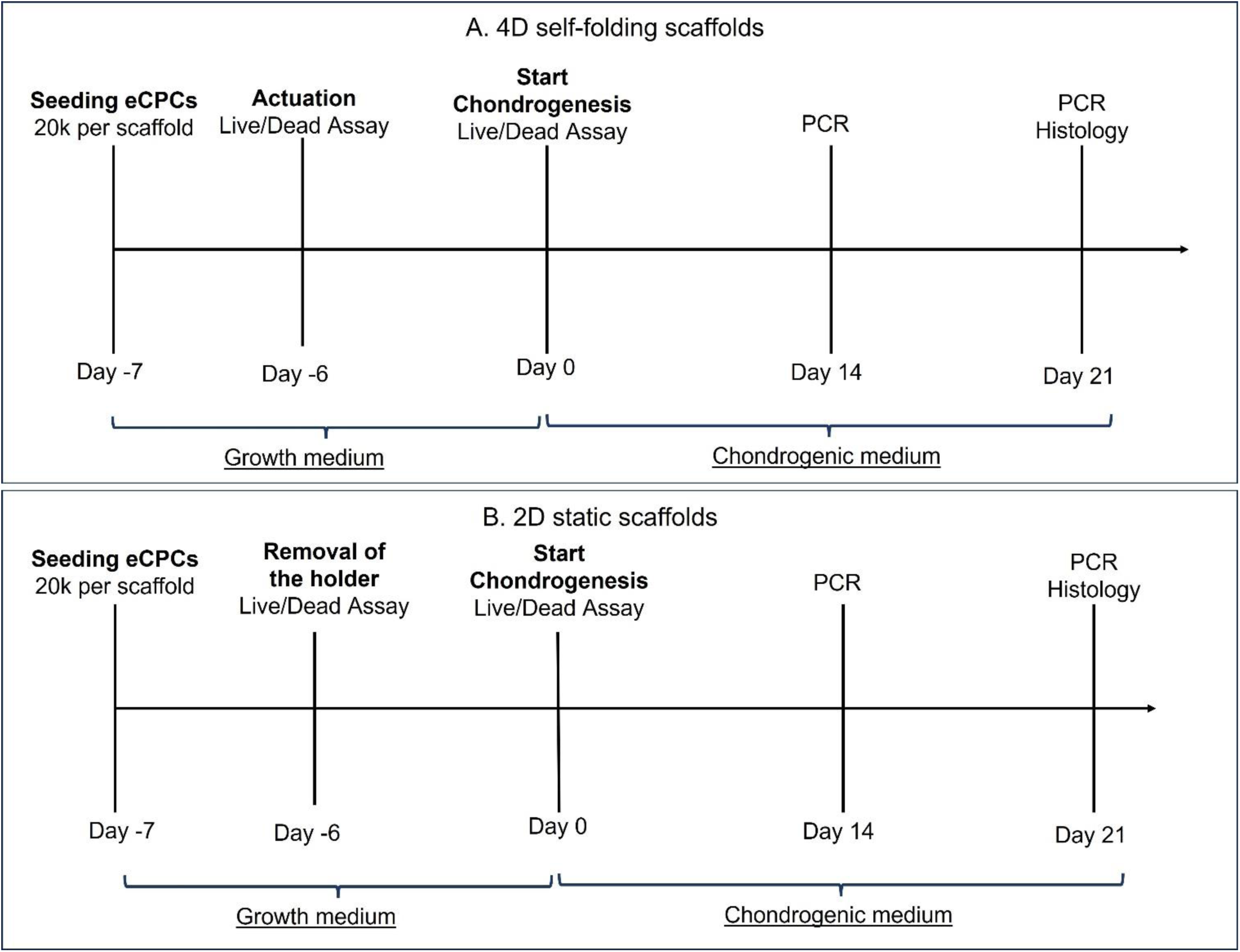
Timeline of the study. The same timeline was used for the 4D self-folding scaffold (A) and for the 2D static control (B).

#### 2.6.1 Viability assays

A LIVE/DEAD^®^ Viability/Cytotoxicity Assay Kit (Molecular ProbesTM) was performed at 24 h, 72 h, and 7 days after seeding on both the 4D and 2D scaffolds to visualize live cells distribution and the presence of dead cells on the scaffold. Briefly, cell-seeded constructs were covered with the Calcein AM/Eth-D-1 (4 μM/8 μM) for 45minutes at room temperature, in the dark, and then observed using a BZX810 All-in-One Microscope (Keyence).

#### 2.6.2 Histological assays

After 21 days of the chondrogenesis, the constructs were fixed in 4% paraformaldehyde in PBS 1X solution, embedded in paraffin and cut in 7μm slices. Hematoxylin and Eosin (H&E) and Alcian Blue staining were used to study cell location and glycosaminoglycans (GAGs) deposition, respectively.

#### 2.6.3 Real-time PCR

Total RNA was extracted using Trizol (Invitrogen) following the manufacturer protocol and purified with the RNeasy Plus mini kit (Qiagen). Reverse transcription was performed using the SuperScript III kit (Invitrogen) with random hexamers primers for the preparation of cDNA. Real-time PCR was performed using the StepOnePlus thermocycler (Applied Biosystems) and SYBR Green Reaction Mix (Applied Biosystems). Expression of chondrogenic genes, SOX-9 (*SOX9*), collagen type II (*COLII*), aggrecan (*ACAN*), of the articular cartilage commitment gene proteoglycan 4 (*PRG4*), and the hypertrophic marker collagen type X (*COLX*) was assessed. Primer sequences, shown in Table 2, were verified on GenBank®. The *18S* gene was used as housekeeping gene. Gene expression fold of change was calculated by the comparative cycle threshold (CT) method, using expression levels at day 0 (*i.e*., 24h after seeding) as reference for the 2^-ΔΔCT^ calculation.

**Table 2:**
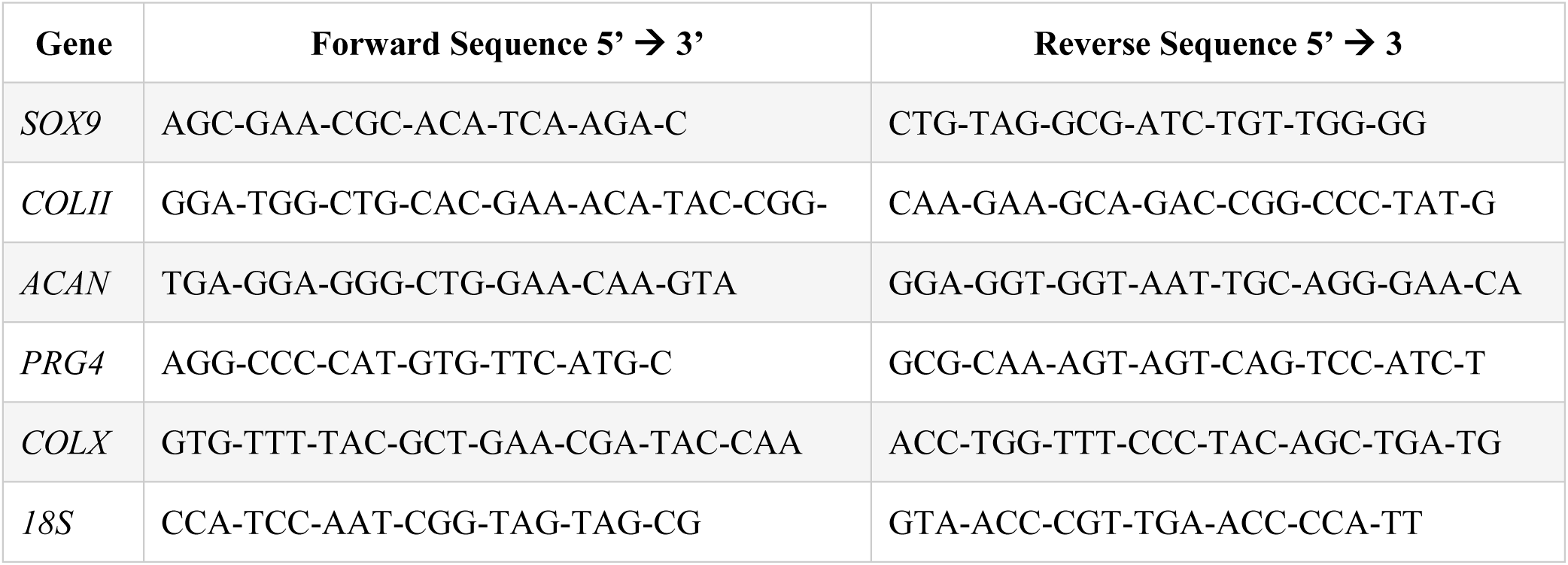
Forward and reverse sequences of the gene primers investigated in the RT-PCR.

### 2.7 Statistical Analysis

GraphPad Prism 7 was used for all statistical comparisons in this work. We used a 1-way ANOVA or a 2-way ANOVA with Tukey’s *post hoc* analysis (multiple comparison) for comparison between groups, with one or two independent variables. A t-test was used when only two groups are compared. In all analysis, the significance thresholds were indicate as *p < 0.05, **p ≤ 0.01, ***p ≤ 0.001 and ****p ≤ 0.0001.

## 3. Results

### 3.1. Materials Characterization

GPTMS-GEL-15 and GPTMS-GEL-5 films were first characterized via swelling tests. The swelling coefficients were analyzed within each material to assess any anisotropy. Statistical analysis revealed that no differences in swelling coefficient between gels in the x and y directions (Figure 5-Ai-ii). Thus, hypothesis (ii) of the analytical model (described in SI1) is verified and a unique swelling coefficient in the x-y plane can be obtained for both solutions. Notably, for both solutions the swelling coefficient is statistically different along z from that on the x-y plane (Figure 5-Ai-ii). The swelling coefficients between the two materials significantly differs in both the x-y plane (Figure 5A-iii, p<0.0001) and along z (Figure 5A-iv, p<0.0001). This inter-material difference in swelling coefficients is the driving force for the folding of the scaffold.

**Figure 5:**
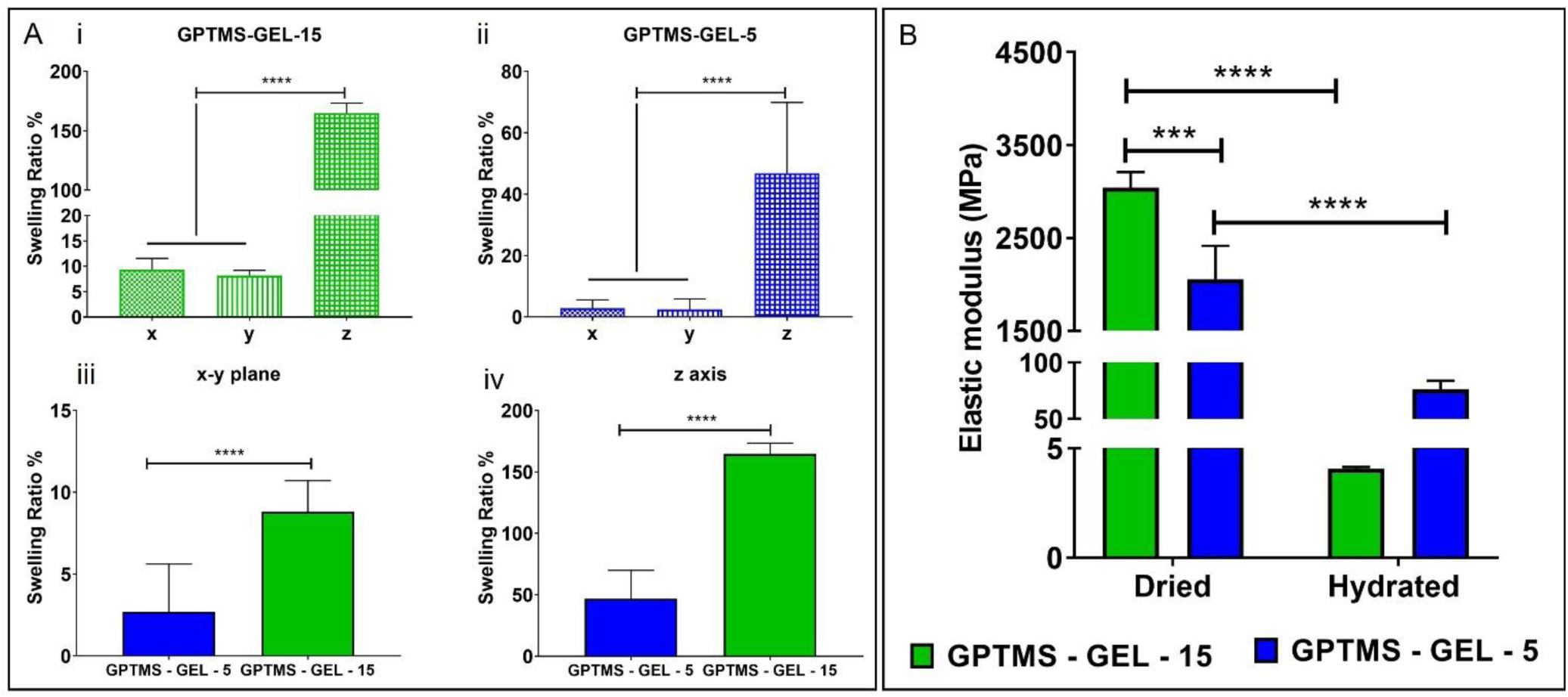
A) Swelling coefficients of the involved materials. i-ii) Swelling coefficient along x, y, and z for each solution. **** p<0.0001. iii-iv) Comparison of the swelling coefficient along the x-y plane (iii) and along the z-axis (iv) inter materials, **** p<0.0001. B) Elastic modulus of the involved materials according to the film state, *** p < 0.001.

Then, the elastic modulus of both materials was evaluated, in dry and hydrated conditions (Figure 5B), which was in the order of GPa in dried state and decreased, as expected, to the order of MPa after hydration for both materials.

The averaged values of swelling coefficients and elastic moduli were used as input for validation of the model.

### 3.2. Characterization of the self-folding scaffolds

Bilayer scaffolds made of GPTMS-GEL were fabricated combining solvent casting and extrusion-based 4D bioprinting. Morphological analysis of the dried film showed a global film thickness between 60 μm and 100 μm, a bioprinted line width of 619 ± 62 μm and a line distance of 1150 ± 115 μm.

When hydrated, the 4D bioprinted scaffold self-folds in approximately 2.5 minutes, maintaining its shape afterwards (Figure 6A). The actuated tubular scaffolds could be easily handled and maintained shape when removed from PBS (Figure 6B). The second layer made of precisely oriented lines provides a complete control over the film folding, thus allowing to obtain two different folding directions determined by the line orientation (Figure 6B). The consequent final tubular structure will have different dimensions in terms of internal diameter, external diameter, thickness, and length depending on the direction of folding, as reported in Table 3.

**Figure 6:**
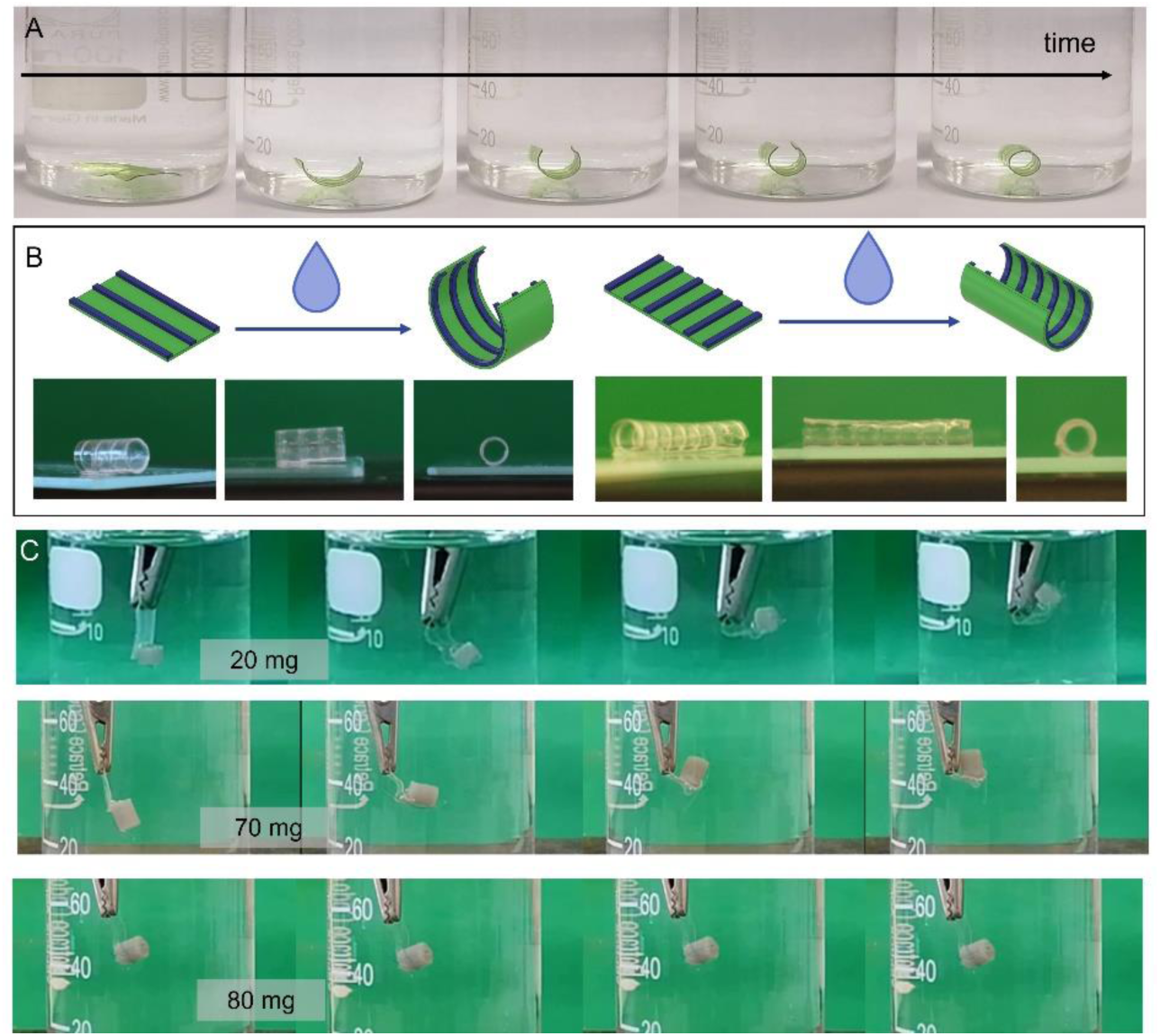
A) Self-folding over time of the scaffolds (10 mm x 5 mm – bioprinted lines parallel to the long side of the scaffold). Food dyes were added in the material only to enhance scaffold visual display). B) The orientation of the lines precisely controls the folding of the 4D bioprinted scaffold (10 mm x 5 mm). C) Experimental evaluation of the force exerted by the scaffold during its self-folding. Up to 70 mg, the scaffold is able to self-fold in time. Differently, when an 80 mg weight is used the folding is not achieved.

**Table 3:**
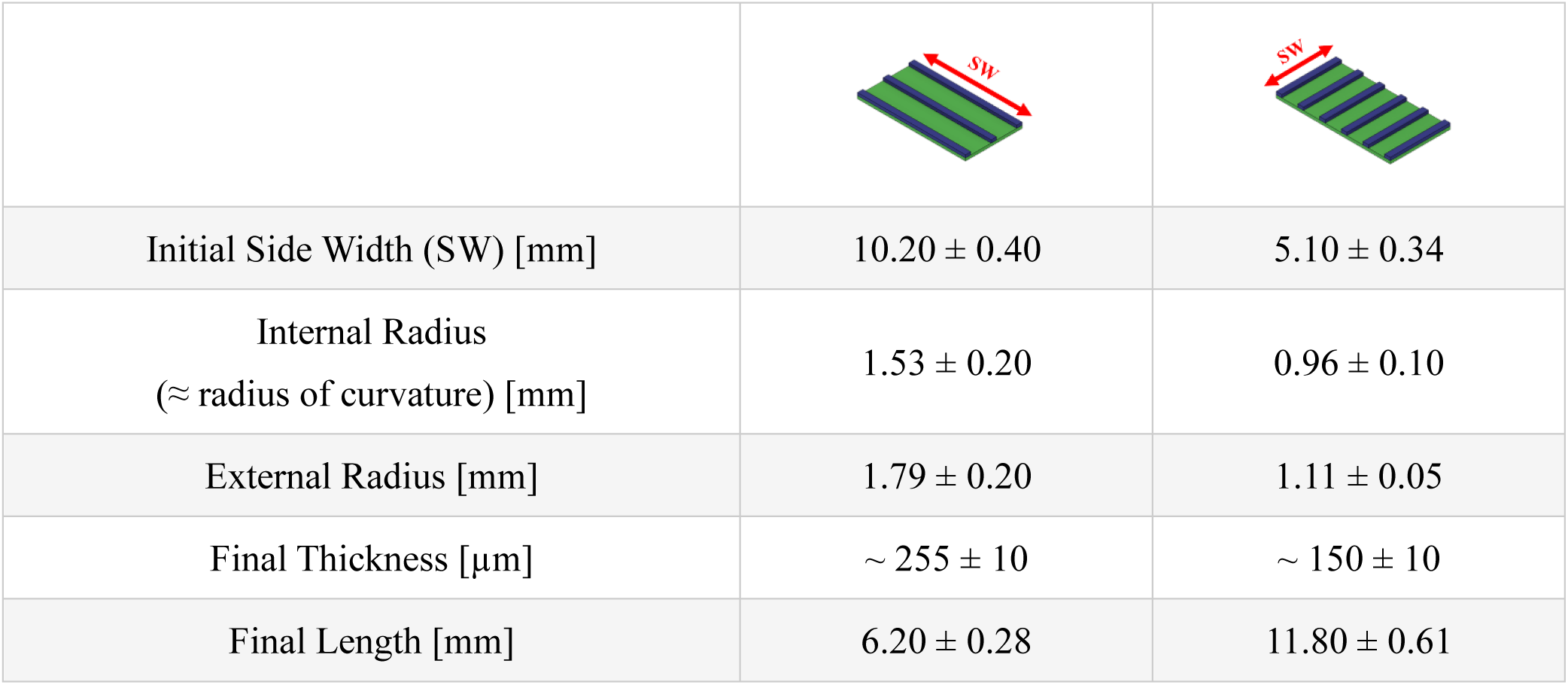
Morphological analysis of the self-folded scaffolds, in relation to the different orientations of the bioprinted lines. Data are reported as mean ± std.

|  |  |  |
| --- | --- | --- |
| Initial Side Width (SW) [mm] | $10.20 \pm 0.40$ | $5.10 \pm 0.34$ |
| Internal Radius<br>(≈ radius of curvature) [mm] | $1.53 \pm 0.20$ | $0.96 \pm 0.10$ |
| External Radius [mm] | $1.79 \pm 0.20$ | $1.11 \pm 0.05$ |
| Final Thickness [$\mu\text{m}$] | $\sim 255 \pm 10$ | $\sim 150 \pm 10$ |
| Final Length [mm] | $6.20 \pm 0.28$ | $11.80 \pm 0.61$ |

The amount of force exerted by the scaffolds during folding was also assessed. The scaffolds were able to self-fold when a weight up to 70 mg was hanged off them (videos of the actuation are shown in SI7.1 and SI7.2), thus resulting in an exerted force of 0.1037 mN (Figure 6C)

### 3.3 An analytical model to predict the scaffolds self-folding

To predict the radius of curvature of the 4D scaffold, we developed an analytical model based on Timoshenko’s bimetal thermostats to relate the radius of curvature of the scaffold to the geometrical parameters and material properties. The analytical model is summarized by Eq. 5

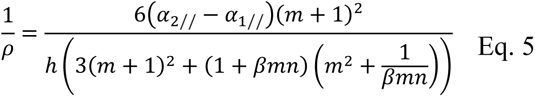

where 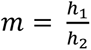 and 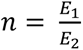. Figure 7 shows how the radius of curvature varies according to the dimensionless numbers m, n, and β that characterize the model.

**Figure 7:**
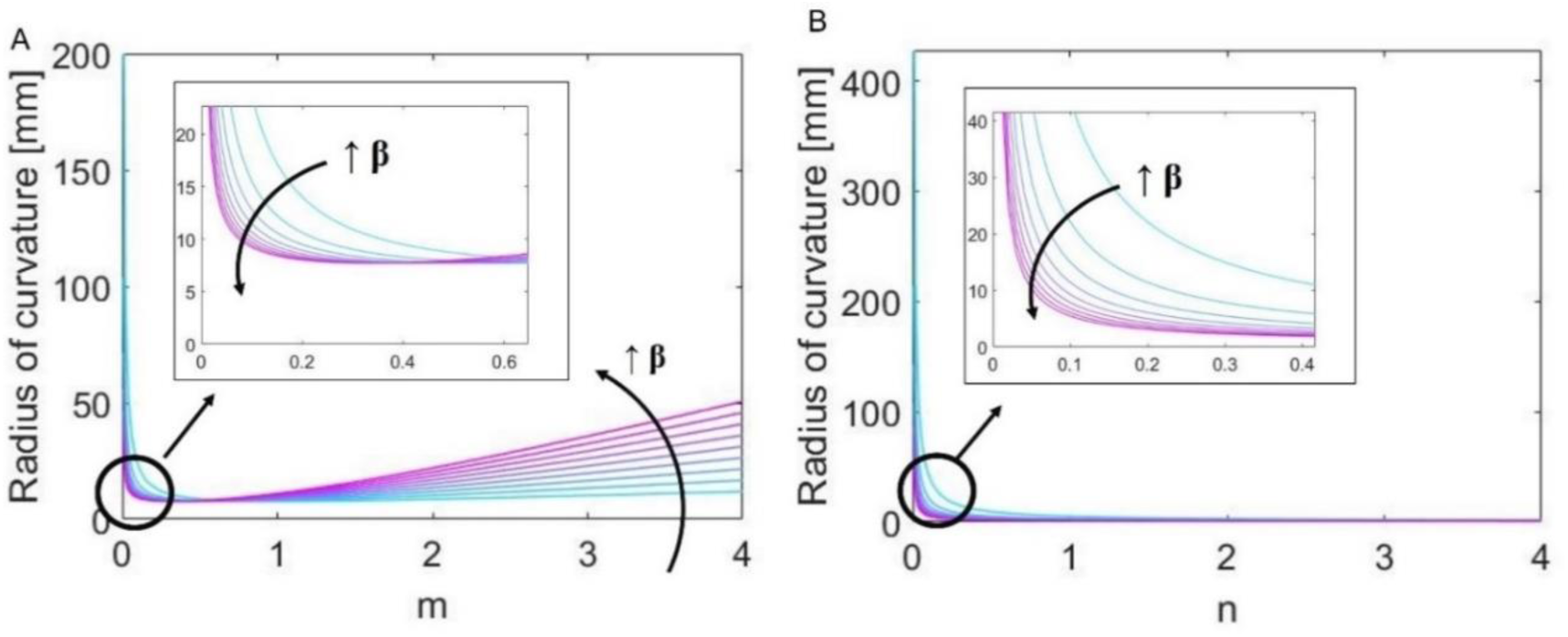
Trend of the radius of curvature predicted by the model according to the dimensionless numbers that characterized it: i) m (i.e., the ratio between the thickness of the two materials); ii) n (i.e., the ratio between the elastic modulus of the two materials); iii) β (i.e., the duty cycle).

### 3.4 4D bioprinted scaffold promotes the proliferation of different airway-related cell sources

To test cell-scaffold interaction, AFs and ECs were seeded on the internal-to-be layer of the scaffolds, whereas eCPCs on the external-to belayer. All cell populations (Figure 8) attached easily to the scaffold in its flat position, proliferated over time (p < 0.0001, two-way ANOVA), and maintained adhesion during the scaffold folding.

**Figure 8:**
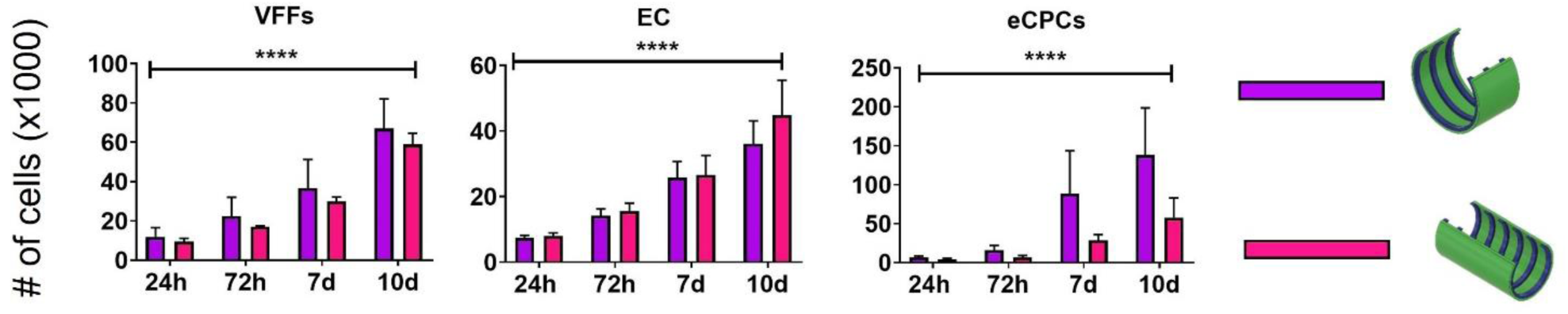
Proliferation assay via Alamar Blue® test performed on the 4D scaffolds, on both folding orientations. Two-way ANOVA revealed a statistically significant differences over time for all cell types. **** p < 0.0001

Higher seeding efficiency was achieved for AFs and ECs, 59.4 ± 2.3 % and 37.5 ± 3.6 %, respectively, with instead a 30.0 ± 1.2 % seeding efficiency for eCPCs(Table 4).

**Table 4:**
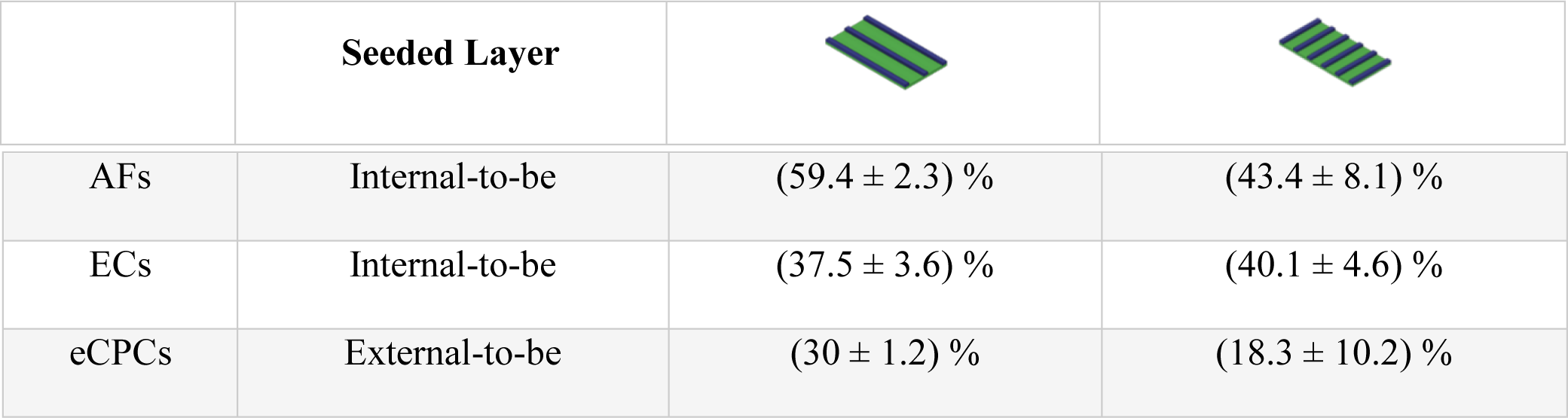
Seeding efficacy for each cell type according to the different bioprinted line orientation. Data are reported as mean ± std.

Since for eCPCs, seeding on the scaffold with bioprinted line parallel to the long side resulted in a higher proliferation rate, (Figure 9, black), we adopted this configuration for the differentiation experiment.

**Figure 9:**
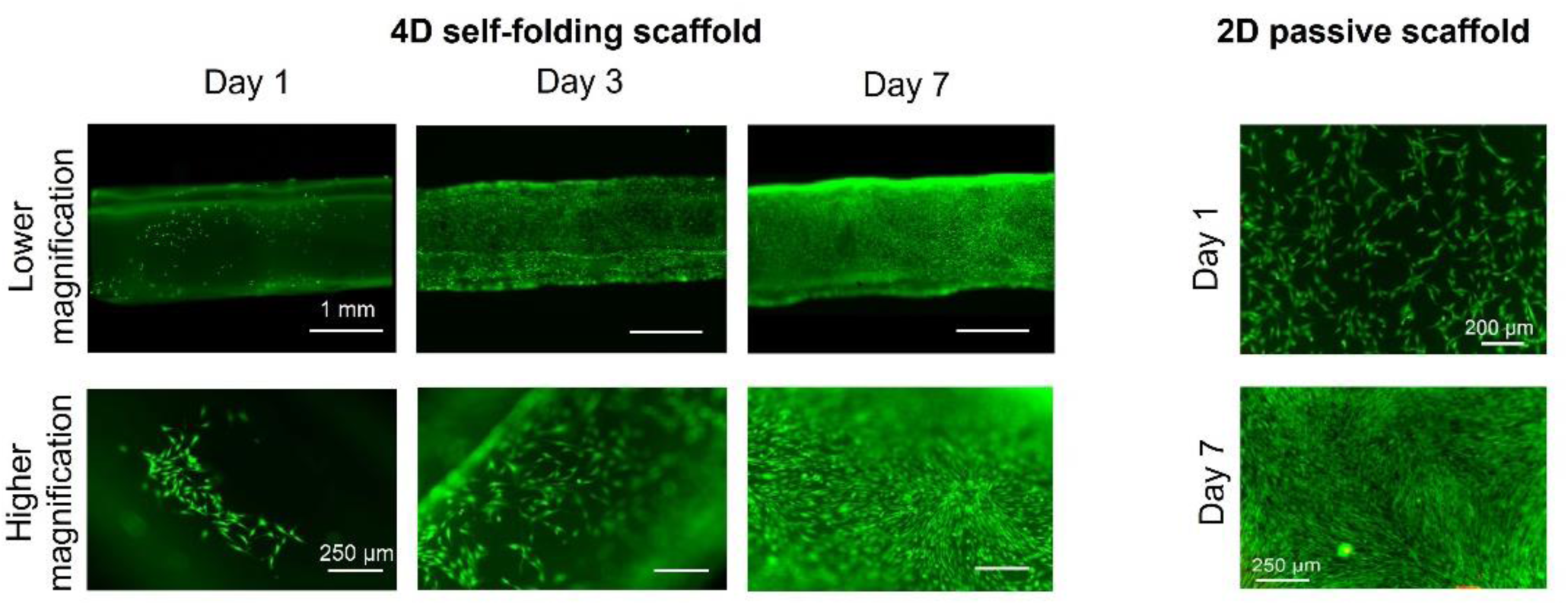
Live/Dead assays over time on the 4D and 2D scaffolds. Cells were able to proliferate and completely cover the scaffold in 7 days in growth media. No morphological differences can be seen between the 4D scaffold and the 2D one at day 7.

### 3.5 4D bioprinted scaffolds promotes the chondrogenesis of the eCPCs

The capability of eCPC to differentiate towards mature chondrocytes on the 4D bioprinted tube was compared with the same cells cultured on 2D static monolayer films.

Scaffolds with lines parallel to the long side were used in this study, since they showed a better cell seeding efficacy (Table 4). eCPCs were seeded on the scaffold in its flat position and, after scaffold actuation (24 hours after the seeding), cells were let free to proliferate for 7 days in growth medium. Live/Dead assays confirmed cells covered the whole scaffold outer surface (Figure 9), and no dead cells can be seen on the scaffolds, confirming its high biocompatibility. Moreover, no morphological differences were visible between cells cultured in the 4D scaffolds versus cells cultured on the corresponding 2D scaffolds of the same composition. Then, cells were cultured in chondrogenic medium for 21 days. Upon visual inspection we observed the deposition over time of white, cartilage-like matrix on the 4D scaffolds suggesting and effective chondrogenic commitment of the eCPCs (Figure 10A). Histologcal staining by hematoxylin and eosin confirmed the persistence of eCPCs on the external layer of the scaffold (Figure 10B-i) whereas Alcian Blue, shown in Figure 10B-ii, stained deep blue further confirming the deposition of cartilaginous matrix.

**Figure 10:**
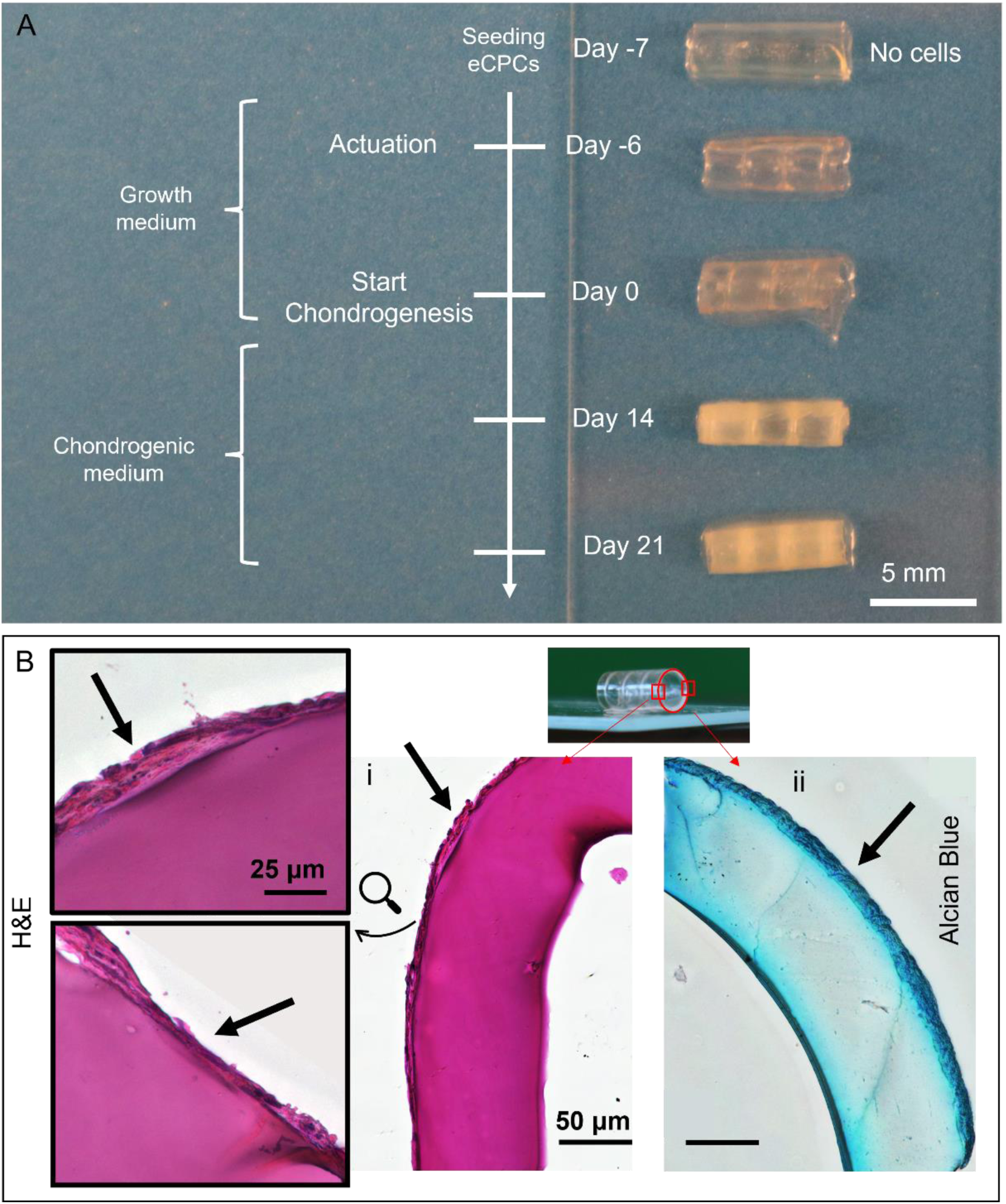
A) Visual inspection of the scaffolds over time, showing the increasing deposition of a white cartilage-like matrix on the scaffold by cells. Scaffolds possess the 2-layer lines parallel to the long side. B) Histological assays performed on the scaffold: i) H&E confirms the presence of an eCPCs layer on the other layer of the scaffold (arrows); ii) Alcian Blue confirms the deposition of GAGs by the eCPCs, and, as a consequence, the formation of a cartilaginous ring (arrow).

Finally, the chondrogenic differentiation of eCPCs was further quantified by RT-PCR, reported in Figure 11, where differentiation on the 4D bioprinted scaffold is in green and on the 2D scaffold in orange. Chondrogenesis is marked by the overexpression of early (*SOX9*) and mature chondrogenic genes (*ACAN* and *COLII*) with statistically significant difference from day 0 (right before culture in chondrogenic medium). Therefore, we can infer that eCPCs on the 4D scaffold formed mature cartilage after 21 days of culture with no hypertrophy (no upregulation of *COLX*). Interestingly, when cells cultured on the 4D scaffolds are compared with the one cultured on the 2D scaffold, marked differences in the gene expression are present. Indeed, at 21 days, cells cultured on the 4D scaffolds (Figure 11, green) overexpressed *SOX9*, *ACAN* and *COLII*, hallmark cartilage genes, to a statistically significant higher extent than cells from the same donors on the 2D scaffolds (Figure 11, orange). Moreover, PRG4, a gene associated to the surface cartilage layer in synovial joints, shows no differences in expression between 2D and 4D scaffolds, suggesting that 4D actuation does not lead to an articular phenotype. Similarly, there is no statistically significant difference in the expression of *COLX* between 2D and 4D.

**Figure 11:**
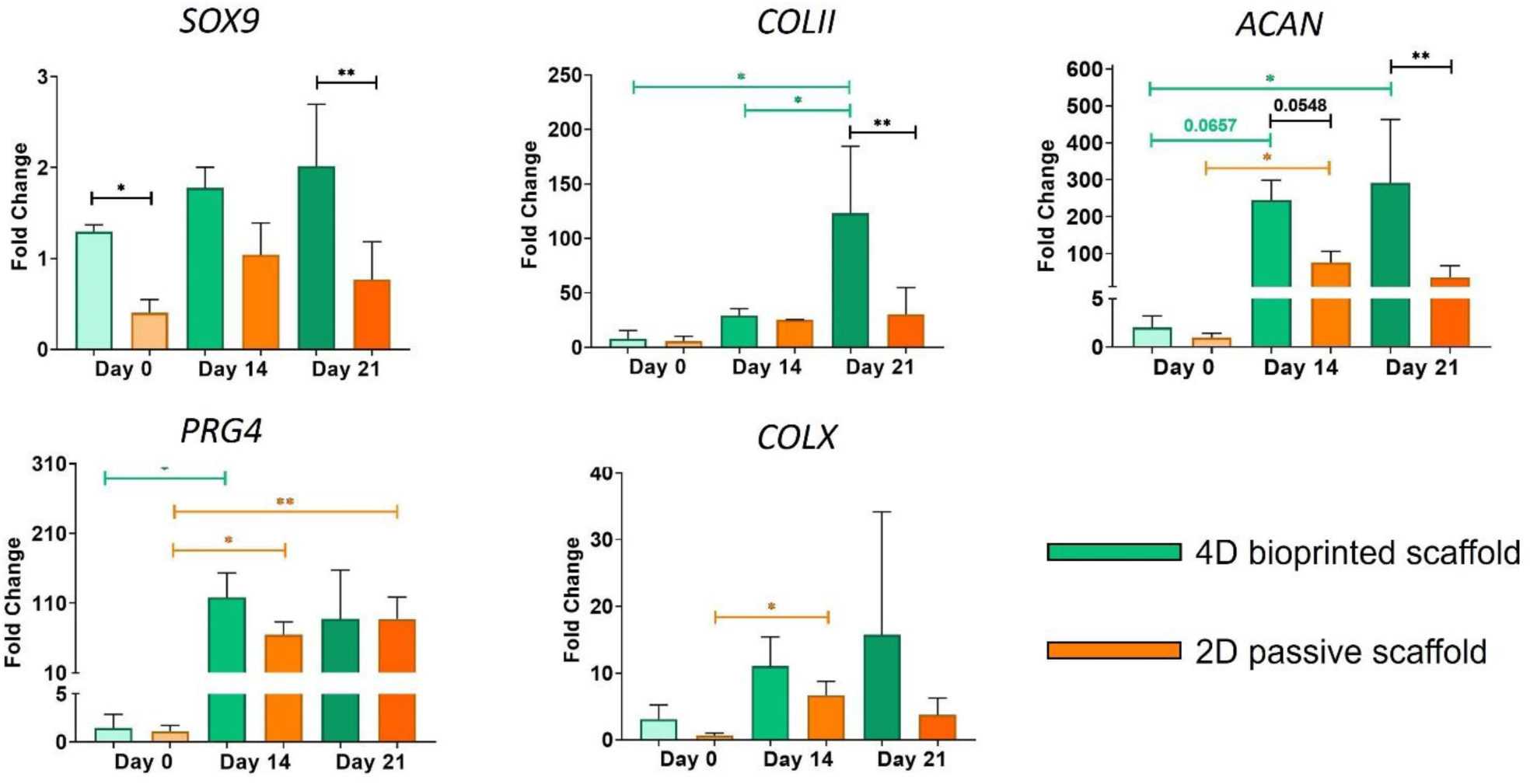
RT-PCR for eCPCs seeded and cultured on the 4D bioprinted scaffolds (green) and on the 2D passive control (orange) for the chondrogenic genes SOX9, COLII, ACAN, and PRG4, and for the hypertrophic gene COLX. * p < 0.05, ** p< 0.01.

## 4. Discussion

In this work, we developed a 4D bioprinted bilayer scaffold able to self-fold upon hydration for the engineering of trachea. The desired self-folding over time was achieved thanks to the differential swelling coefficient in aqueous solution of the two layers that compose the scaffold (Figure 5.A). Thus, when the scaffold is dipped into a water-based solution the layers’ swelling mismatch creates a deformation gradient that drives the scaffold folding (Figure 6.A). This is a well-described phenomenon in literature that was applied in several studies, employing both synthetic [27], [28] and natural polymers [29]. For instance, Aspite *et al.* [30] electrospun a bilayer scaffold made of PCL and poly(N-isopropylacrylamide) (PNIPAM) that could self-fold in aqueous environment at 37°C, due to the PNIPAM shrinkage. Differently, Lee *et al.* [31] created bilayer films based on alginate and gelMA with a gradient in crosslinking that led to a difference in layer swelling rate, thus driving the structural deformation.

In this work, we selected GPTMS-crosslinked gelatin as bulk material for both layers because of its high biocompatibility and good cell affinity, as we shown in [24], [32], [33]. Further, to have both layers made with the same bulk material guarantees a covalent chemical bond between the layers, thus preventing delamination, for the whole length of the experiment of 28 days of cell culture. Adjusting gelatin concentration and GPTMS content, we were able to tune the differential swelling behaviour of the (Figure 5.A). In fact, our data showed that the increase in gelatin content leads to an increase in the swelling coefficient due to the higher amount of water-absorbing polymer. On the other hand, the increase in GPTMS leads to a decrease in the swelling coefficient due to the higher degree of crosslinking.

Other than swelling coefficients, there are several variables, both related to the materials properties (*e.g.*, elastic modulus) and to the geometry of the scaffold (*e.g.*, layer thicknesses and number of bioprinted lines), that influence the folding of the scaffold. Thus, to precisely predict the radius of curvature, we developed a forward analytical model, summarized in Eq. 5, based on the Timoshenko’s bilayer thermostats [26]. Our model showed that the radius of curvature of the scaffold is directly proportional to its total thickness and inversely proportional to the difference between the expansion coefficients along the x-y plane. Similarly to Timoshenko’s approach, our model showed a dependence on the ratio of the thicknesses of the two layers (m) and on the ratio of the elastic moduli of the two materials (n). Differently from Timoshenko’s model, the β term appears, which takes into account the number of bioprinted lines (N), their width (D), and the width of the entire scaffold (W), thus incorporating the periodical geometric features that develop in the plane, which are not present in Timoshenko’s one-dimensional model. Moreover, according to our model the side width (SW) does not affect the final radius of curvature of the structure. However, the SW (that represent the circumference of the actuated scaffold) must to be carefully tuned in the design phase, so that, after the actuation, depending on the application either a close tube, or a multiple folded (*e.g.*, cochlear engineering), or a partially curved structure (*e.g.*, condyle engineering) can be obtained. The model was then validated according to three key parameters of interest: i) the total thickness of the structure, that Timoshenko identified as the parameter that most influences the curvature of the structure; ii) β, that takes into account the bioprinted pattern, and iii) the external dimensions of the film that are involved in the scalability of the structure. It is important to stress out that our model is a general-purpose model that can be exploited to describe the folding behavior of any bilayer planar structure under deformation mismatch, and it is not constrained to a specific fabrication approach, the selected materials, or the exploited stimulus, thus it could be applied to several other applications besides the one presented in this work.

For the fabrication of the scaffold, we exploited the 4D bioprinting approach that leverages additive manufacturing of active and passive materials to create active scaffolds that exhibit an environmental-triggered predefined shape transformation. 4D bioprinting can be seen as the use of 4D printing (*i.e.*, an innovative fabrication approach that adds time, the 4^th^ dimension, to AM, introduced by Dr. Skylar Tibbis in 2013 [34]) applied to tissue engineering for the fabrication of dynamic scaffolds. In 4D bioprinting the selection of the appropriate biomaterials and related stimulus is crucial since they should simultaneously lead to the desired actuation and be biocompatible. This last condition drastically restricts the plethora of usable biomaterials and stimuli, thus limiting the advancement of this fabrication approach. Here, both the biomaterials (*i.e.,* GPTMS-GEL based solution) and the stimulus (*i.e.,* hydration) are highly biocompatible and safe to use with different cell types.

Overall, 4D bioprinting constructs possess several advantages over static bioprinting, such as facilitating the seeding process and the possibility to adapt to complex anatomical structures [15], [19]. Here, the 4D bioprinting approach specifically allows to: i) facilitate the seeding of different cells populations on either side of the structure in its initial flat position; ii) obtain a tubular structure that resembles the morphology of the native trachea, as the geometric form of a tissue/organ is inherently linked with its biological functions at all scales [35]; iii) provide mechanical stimuli to cultured cells leading to improved cartilaginous maturation of eCPCs, as discussed in details later on.

In 4D bioprinting, AM acts an enabling technology that allows the deposition of one or multiple materials with a precise and predefined spatial organization with negligible complexity constraints. Here, we adopted extrusion-based bioprinting as the AM technology to deposit the GEL-GPTMS-5 biomaterial ink with a precise line patterned geometry to finely control the folding direction of the film (Figure 6B). Indeed, the folding direction is dictated by the bioprinted line orientation, so the axis of the obtained tube is perpendicular to the bioprinted lines. A similar phenomenon was already observed by Yang *et al.* [29], where the self-folding of a gelatin-based film was controlled by the direction of equally distributed grooves on the film surface, so that the folding direction was perpendicular to the direction of the grooves. Mechanically, the presence of the bioprinted lines (as well as the grooves in Yang’s study) drives the direction of the folding since the lines width (D) is for all intents and purposes negligible if compared with the structure width (W). Thus, the folding radius of curvature along the y direction (Figure 3A) is predominant over the other directions. In other words, each line acts as a Timoshenko beam driving the folding along its length with negligible effect on the other directions.

Interestingly, the developed 4D bioprinting fabrication approach alongside with the selected materials resulted in the fabrication of scalable scaffolds that could be designed to achieve a wide range of diameters in their final configuration. In fact, exploiting the developed analytical model and designing the initial scaffold geometry accordingly, we obtained a tubular structure with a diameter ranging from 1.5 mm to 15 mm (Figure SI3). This would allow the use of the develop 4D bioprinted tracheal construct for *in vivo* models of different sizes (*e.g*., mouse, rabbit, pig [36], [37]) and potentially also in clinical studies of patients of different ages (e.g., newborn, teenager, adult [38], [39]). Moreover, the easy scalability of the model would allow to engineer structures for other tubular anatomical areas, such as vasculature or nerves.

In this study, for the biological evaluation of the scaffold, we used the 10 mm x 5 mm scaffold (diameter between 2 mm and 3 mm) to have an easy-to-handle scaffold that could fit well into a 12 well-plate and has the potential in the future to be deployed in an animal model. First, we assessed the possibility of seeding and culturing the different cell populations of the airway, with the ultimate aim to fabricate a hierarchical graded scaffold able that could mimic the stratified morphology of the native trachea. Immortalized human airway fibroblasts [40], [41] were used to mimic the connective tissue on the inner side of the trachea and were seeded in the internal-to-be-layer of the scaffold. Similarly, immortalized non-tumorigenic human lung epithelial cells were seeded in the internal-to-be-layer of the scaffold to mimic the tracheal epithelium. Then, human ear cartilage progenitor cells, harvested form paediatric patient surgical waste samples were used to mimic the cartilaginous tracheal rings and were seeded in the external-to-be layer of the scaffold. All cell populations successfully attached to the scaffold in its flat position, remained adhered during the scaffold folding, that occurs 24 h after seeding, and proliferated on the scaffold, as evidenced by a statistically different cell number over time (Figure 8). In terms of seeding efficacy, after the actuation the eCPCs reached the lowest value, of approximately 30%. This could be due to the fact that AFs and ECs being cultured on the inner side of the scaffold, could be more protected from the stresses associated to scaffold manipulation and medium change. Overall, the seeding efficacy could be considered satisfactory, and the gap between the number of cells seeded and cells counted after 24h can be accounted for by different degrees of cell attachment to the holder, cell detachment during scaffold folding, scaffold manipulation, and medium change. Notably, we did not observe any morphological change in the scaffold geometry, as well as any de-folding of the structure during and after cell culture. This highlights the mechanical stability of the developed structure, both in terms of fabrication approach and material selection. A similar result was achieved by Díaz-Payno *et al.* [16], who developed a self-folding scaffold based on alginate and hyaluronic acid for the engineering of curved cartilage (such as the ear or the condyle). The authors showed that after 28 days in culture, the curvature (that for their study is around 0.15 mm and does not lead to the complete closure of the structure) was still clearly present and did not change over time.

Finally, we tested the possibility of realizing a single cartilage ring, arguably the hardest tracheal component tissue to engineer, by culturing eCPCs on the self-folding scaffold. eCPCs are a small but highly proliferative cell population, characterized by rapid self-renewal, that resides within ear cartilage. Recently we showed that eCPCs are able to proliferate faster than chondrocytes and mesenchymal stem cells, thus being a promising cell source for cartilage engineering [42]. When comparing by real-time PCR the gene expression of the eCPC cultured on the 4D bioprinted scaffold versus that of the eCPCs on the 2D passive scaffolds, we observed notable (Figure 11). Indeed, at 21 days of differentiation ***SOX9***, ***ACAN*,** and ***COLII*** were significantly more overexpressed on the 4D scaffolds compared to the 2D ones. Moreover, there were no differences between 4D and 2D in the expression of PRG4, a gene associated to the surface layer of articular cartilage, thus suggesting that the 4D scaffold while promoting chondrogenesis, does not specifically skew towards an articular phenotype. Importantly, the absence of any difference in the expression of ***COLX*** suggested that cells are not driven towards hypertrophy, suggesting the establishment of a stable chondral phenotype on the 4D scaffolds. Moreover, no differences were observed between the 2D and the 4D scaffolds in terms of cell density or cell confinement (Figure 8), that has been showed in literature to positively influence chondrogenesis [43], [44]. Together these results strongly suggest that culturing eCPCs on the 4D bioprinted self-folding scaffolds improves chondrogenesis, specifically for airway engineering (no differences in the expression of *PRG4* and *COLX*). This implies that eCPCs positively respond to the scaffold folding and its final curvature. Different works have shown that cells can discriminate between planar, convex, and concave surfaces and that can be influenced by curvature, when the radius is in the milli- or micro-range [35], [45], [46]. For example, Lee *et al.* [47] showed that NIH-3T3 fibroblasts were sensitive to the curvature of glass balls with a diameter up to 2 mm. Furthermore, Werner *et al.* [48] showed that culturing human mesenchymal stem cells on convex surfaces (diameters ranging from 0.25 to 0.75 mm) enhances their osteogenic commitment, but decreases their migratory capacity. The authors hypothesized that this could be caused by the convex surfaces leading to a full-contact of the cells to the substrate and, as a consequence, causing the flattering of the cell nuclei that enhances osteogenic differentiation.

In this context, to the best of our knowledge, our study showed for the first time how the formation of healthy cartilage for the airway could be enhanched by the 4D geometry of a scaffold. Notably, previous studies on self-folding scaffold for cartilage tissue engineering did not show or analyse any quantitative differences versus a static control [16], [19], [23], [31]. In our study, we evaluated how cells are subjected to a strain, equal to approximately 6.47% (Eq.SI15), exerted by the folding scaffold due to its curvature. Thus, we can speculate that, by exercising this mechanical force, the 4D bioprinted self-folded scaffold regulates cartilage formation, possibly by mimicking the action of smooth muscle cell progenitors. Indeed, these cells, during trachea morphogenesis in the embryo, provide the biomechanical stress-strain configuration that regulates and promotes trachea tubular shape and cartilage formation [36], [49].

## 6. Conclusion

In this study, we exploited the 4D bioprinting approach to fabricate a self-folding bilayer scaffold for trachea engineering. The scaffold is able to change its shape upon hydration, reaching a tubular structure with a programmable final diameter, determined by its geometrical features and material properties. Biologically, we firstly analyzed the ability of different cell sources of airway interest to attach to the scaffold in its initial flat position, to maintain their adhesion upon actuation, and proliferate over the scaffold in its final folded position. eCPCs were able to differentiate towards mature cartilage and, interestingly, our data reveiled how the cells responded to the scaffold folding and final curvature with an increase in the upregulation of pro-chondorgenic genes associate with a stable and robust cartilage phenotype. We speculated that upon scaffold folding, the cells sensed a mechanical strain resembling that generated by smooth muscle progenitors in the embryonic tracheal development which leads to enhanced tracheal cartilage ring formation. This result represents an important milestone for 4D bioprinting, quantitativelly showing that shape-morphing scaffolds in the millimeter scale can promote cells activity and differentiation.

## Acknowledgements and Fundings

Airway Fibroblasts were kindly provided by Dr. Susan L. Thibeault, from the University of Wisconsin, Madison. The authors are thankful to Dr. Soheila Ali Akbari Ghavimi for eCPCs extraction. The authors would like to thank Carlotta Salvatori, University of Pisa, for performing preliminary bioprinting tests.

I.C., A.E., G.V., C.D.M. acknowledge the support of the Crosslab Additive Manufacturing of the Department of Information Engineering of the University of Pisa and the support of the FoReLab of the Department of Information Engineering of the University of Pisa.

We acknowledge funding support by the National Heart, Lung, and Blood Institute of the National Institute of Health R21HL159521 and 1R56HL164536-01 (RG), the Children’s Hospital of Philadelphia Research Institute (RG), the Frontier Program in Airway Disorders of the Children’s Hospital of Philadelphia (RG).

