## Supplementary Information for "4D bioprinted self-folding scaffolds enhance cartilage formation in the engineering of trachea"

### *SI1. Analytical Model Hypothesis and Derivation*

The model is based on the following hypothesis:

i. where both layers are present, the geometry can be considered as a bilayer beam, that behaves according to Timoshenko's model. This hypothesis is verified if the width of the lines (D) is negligible compared to the side width (SW);

ii. where only the solid layer is present, it expands in all directions without interacting with the bioprinted upper layer, and consequently without generating the mismatch that drives the folding. This hypothesis is verified if the individual materials exhibit isotropic expansion in the x-y plane;

iii. the deformation given by expansion is not negligible. The geometrical input parameters are considered as already affected by the expansion (*i.e.*, $h_1$, $h_2$, D, and W measured at dry state multiplied by the appropriate expansion coefficient, Eq. SI12, SI13, SI14).

Given hypotheses *i* and *ii* only the internal forces parallel to the lines and the torques relative to them need to be considered. Since no external forces are present, the resultant of internal forces (P) and torques (m) must be zero (Eq. SI1 and SI2).

$$\sum_{1}^{N} \boldsymbol{P_i} = \boldsymbol{P_2} = \boldsymbol{P} \qquad \text{Eq. S1}$$

$$\boldsymbol{P}\frac{\boldsymbol{h}}{\boldsymbol{2}} = \boldsymbol{m_2} + \sum_{1}^{N} \boldsymbol{m_i} \qquad \text{Eq. SI2}$$

Since the printed lines have the same dimensions and are equally spaced, the $P_i$ and $m_i$ are all equal, thus Eq. SI3 can be derived.

$$\sum_{1}^{N} \boldsymbol{P_i} = \boldsymbol{NP_1} \text{ - } \sum_{1}^{N} \boldsymbol{m_i} = \boldsymbol{Nm_1} \qquad \text{Eq. SI3}$$

Combining Eq. SI3 into Eq. SI1 and Eq. SI2, Eq. SI4 and Eq. SI5 can be obtained

$$\boldsymbol{NP_1} = \boldsymbol{P_2} = \boldsymbol{P} \qquad \text{Eq. SI4}$$

$$\boldsymbol{P}\frac{\boldsymbol{h}}{\boldsymbol{2}} = \boldsymbol{m_2} + \boldsymbol{Nm_1} \qquad \text{Eq. SI5}$$

Then, considering the involved material as linear elastic, the torques can be written as in Eq. SI6 for both layers, where $I_1$ and $I_2$ are expressed in Eq. SI7.

$$\boldsymbol{m_1} = \frac{\boldsymbol{E_1 I_1}}{\boldsymbol{\rho}} \qquad \text{Eq. SI6.1}$$

$$\boldsymbol{m_2} = \frac{\boldsymbol{E_2 I_2}}{\boldsymbol{\rho}} \qquad \text{Eq. SI6.2}$$

$$\boldsymbol{I_1} = \frac{\boldsymbol{D{h_1}^3}}{\boldsymbol{12}} \qquad \text{Eq. SI7.1}$$

$$\boldsymbol{I_2} = \frac{\boldsymbol{W{h_2}^3}}{\boldsymbol{12}} \qquad \text{Eq. SI7.2}$$

Including Eq. SI6 in Eq. SI5, Eq. SI8 can be obtained

$$\boldsymbol{P}\frac{\boldsymbol{h}}{\boldsymbol{2}} = \frac{\boldsymbol{NE_1 I_1} + \boldsymbol{E_2 I_2}}{\boldsymbol{\rho}} \qquad \text{Equation SI8}$$

Since the deformation at the interface between the layers must be equal for continuity, Eq. SI9 can be obtained

$$\alpha_{1//} - \frac{P1}{E_1 h_1 DN} - \frac{h_1}{2\rho} = \alpha_{2//} + \frac{P2}{E_2 h_2 W} + \frac{h_2}{2\rho} \quad \text{Eq. SI9}$$

The first term of Eq. SI9 is the deformations on the x-y plane due to the swelling, the second term is the deformation due to internal forces calculated according to Hooke's laws, and the last term is the deformation due to the folding. Then, including Eq. SI9 in Eq. SI8, Eq. SI10 that relates the radius of curvature (ρ) with the parameters of interest, can be obtained.

$$\frac{1}{\rho} = \frac{\alpha_{2//} - \alpha_{1//}}{\frac{h}{2} + \frac{2(E_1 I_1 + E_2 I_2)}{h}\left(\frac{1}{E_1 h_1 ND} + \frac{1}{E_2 h_2 W}\right)} \quad \text{Eq. SI10}$$

Including Equation S7, defining the auxiliar adimensional numbers $m = \frac{h_1}{h_2}$ and $n = \frac{E_1}{E_2}$; and including the duty cycle β, Equation S11

$$\frac{1}{\rho} = \frac{6(\alpha_{2//} - \alpha_{1//})(m+1)^2}{h\left(3(m+1)^2 + (1+\beta mn)\left(m^2 + \frac{1}{\beta mn}\right)\right)} \quad \text{Eq. SI11}$$

It is important to stress out that the values of the geometric parameters in Eq. SI11 are considered after the expansion of the materials, in accordance with hypothesis *iii*. Briefly, $h_1$ and $h_2$ were obtained considering the expansion in the z axis of involved material, respectively (Eq. SI12). Differently, since D is influenced by the expansion of both GEL-GPTMS-5 and GEL-GPTMS-15, the value of D after expansion is calculated employing a Voigt model of the two materials (Eq. SI13). Finally, the expansion of W is calculated as the sum of the expansion of each line (D) multiplied by the number of line (N) and the expansion of the solid layer interposing the printed lines (Eq. SI14).

$$h_1 = h_{1\_before\_actuation}(1 + \alpha_{1per}) \quad \text{Eq. SI12.1}$$

$$h_2 = h_{2\_beforeactuation}(1 + \alpha_{2per}) \quad \text{Eq. SI12.2}$$

$$D = D_{before\_actuation}\left(1 + \frac{\alpha_{1par} E_1 + \alpha_{2par} E_2}{E_1 + E_2}\right) \quad \text{Eq. SI13}$$

$$\boldsymbol{W} = \boldsymbol{ND} + (\boldsymbol{W}_{\boldsymbol{before\_actuation}} - \boldsymbol{ND}_{\boldsymbol{before\_actuation}})(\mathbf{1} + \boldsymbol{\alpha}_{\mathbf{2}\boldsymbol{par}}) \quad \text{Eq. SI14}$$

### *SI2. Analytical Model Validation*

#### *SI2.1 Rationale*

The parameters that influence the radius of curvature can be based on the geometry of the scaffold or on the involved materials. Since scaffolds are made of specific materials that has already been used for several biological application [SI1-SI5], we decided to do not include the material property in the parametric validation of the model. In consequence, the parameter “m” that depends on the elastic module is considered constant in the validation.

Differently, geometrical parameters, that affect the radius of curvature, were used in the parametric validation.

The validation of the model is based on comparing the predicted radius of curvature and the effective radius of curvature measured on the experimental samples, in function of the identified parameters of interest: i) the total thickness of the structure (h); ii) the duty cycle (β); iii) the total dimension of the scaffold (SW, W).

For each validation, the parameter of interest varies in a specific range of values (limited by the fabrication technique), reported in Table SI1. The other geometrical parameters are maintained constant, equal to the average value of the real samples Table SI1. The material properties used in the validation model are reported in Table SI1, in the main text.

| Validation parameter | Constant parameter | Variable parameter | Measured parameter |
|---|---|---|---|
| Total thickness | β (0.50 ± 0.09)<br>h1 (33.1 ± 14.1μm)<br>SW (5.1 ± 0.3mm)<br>W (10.2 ± 0.4mm) | h2:<br>from 55 to 160 μm<br>5 μm step | ρ |
| Duty cycle | h1 (26.25 ± 11.87 mm)<br>h2 (60.00 ± 4.29 mm)<br>SW (5.1 ± 0.3mm)<br>W (10.2 ± 0.4mm) | β:<br>0.163<br>0.115<br>0.089<br>0.072<br>0.061 | ρ |

| | | | |
|---|---|---|---|
| | | 0.053<br>0.046<br>0.04<br>0.037 | |
| Scaffold Dimension | β (0.50 ± 0.09)<br>h1 (33.1 ± 14.1µm) | SW, W and $h_2$ as:<br>10 × 20 × 0.08,<br>10 × 20 × 0.13,<br>20 × 10 × 0.08,<br>20 × 10 × 0.13,<br>20 × 20 × 0.08<br>20 × 20 × 0.13 | ρ |

*Table S1: Parameter values combination for each validation experiments.*

#### *SI2.2 Sample fabrication, actuation and characterization*

The samples for the validation in function of the total thickness and scalability were fabricated as described in 2.2 and actuated as described in 2.3.

For the validation in function of the duty cycle, a different value of D was needed to achieve a significant variation of the parameter of interest. For this reason, a needle of 160µm was used to fabricate the scaffolds (printing parameters: print speed = 10 mm/s, needle diameter =0.16 mm, layer height = 0.05 mm, volumetric flow = 0.08 $mm^3/s$). Moreover, different values of duty cycle were obtained by printing parallel lines at a da distance of 2, 3, 5, 6 and 8 mm.

After the actuation, for each sample the radius of curvature was measured by image analysis via ImageJ.

#### *SI2.3 Model Validation according to the total thickness of the constructs*

Figure S1 shows the radius of curvature plotted as a function of the total thickness. Theorical values (continuous green line) predicted by the model are calculated using the average value of measured material and geometrical parameters (*i.e.*, Young's modulus, expansion coefficients, film thickness, lines thickness) with a confidence interval (CI) (dashed green lines) obtained considering the measured values as a t-student distribution. Then, for each experimental samples, a dot is plotted in the graph at the corresponding total thickness and radius of curvature values. Then, radius of curvature of the experimental samples possessing the same total thickness were averaged obtaining a mean curve (continuous magenta line) and two STD curves (dashed magenta lines). When, the total thickness of the testing sample is between 55 and 105µm the predicted value is comprised between the STD of the experimental values. Thus, it can be concluded that the model in this total thickness region the model is able to predict the curvature of the sample in this range of thickness.

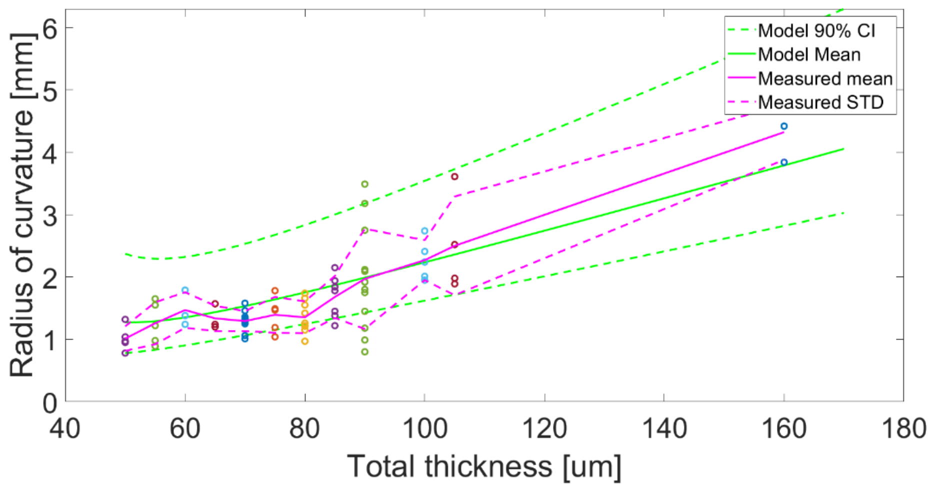


Figure SI1: Validation of the analytical model. Green line indicates the predictive values from the analytical model, whereas the dashed green lines indicate the model 90% confidence interval (CI). Dots represents the individual experimental measurements for each thickness. Magenta lines represent the average and std (dashed line) of the experimental measures.

#### *SI2.4 Model Validation according to the duty cycle*

In Figure SI2, the radius of curvature is plotted as a function of β. Testing samples with different thicknesses are merged. The measured radii of curvature in function of the parameter β are compared with the values predicted by the model. In this case, the number of parameters that influence the variability is increased in confront of the previous validation, for this reason identify a confidence interval is not practicable. Instead of a CI we used the mean value predicted for the higher thickness measured and the lower thickness that were measured. Since measured values followed are between the range of predicted values, we can assume that the model is able to predict the radius of curvature of the structure when β is comprised between 0.05 and 0.2.

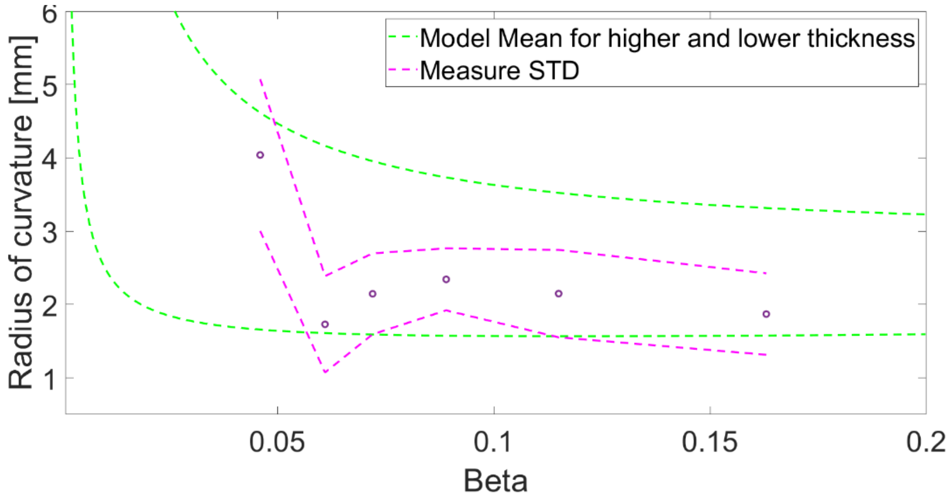


Figure SI2: Second validation of the analytical model according to β. The dashed green line indicates the higher and lower predictive values from the analytical model. Dots represent the mean of experimental measurements for each beta value. The magenta dashed lines represent the std of the experimental measures. …

*SI2.5 Model Validation according to the total dimension of the scaffold*

In Figure SI3, the radius of curvature is plotted as a function of the total thickness. Green continuous lines and green dashed lines are the same as in S2.3, representing the predicted value and its CI. Blue, red, and black dots are testing samples with W and SW respectively equal to 20 x 10 mm, 10 x 20 mm and 20 x 20 mm, with a theorical thickness equal to 80 and 130 µm. Due to fabrication variability the thickness is spread around those value, forming a cluster of dots. For each theorical thickness, the STD of thickness and the radius of curvature is reported as dashed magenta lines. Except for few samples, experimental values fall into the 90% CI region of the model. Moreover, the STD of the predicted values falls into the STD of the testing samples. This suggests that the model is able to predict the radius of curvature of the structure regardless its global dimension. This is extremely useful for the scalability of the model, described in SI3.

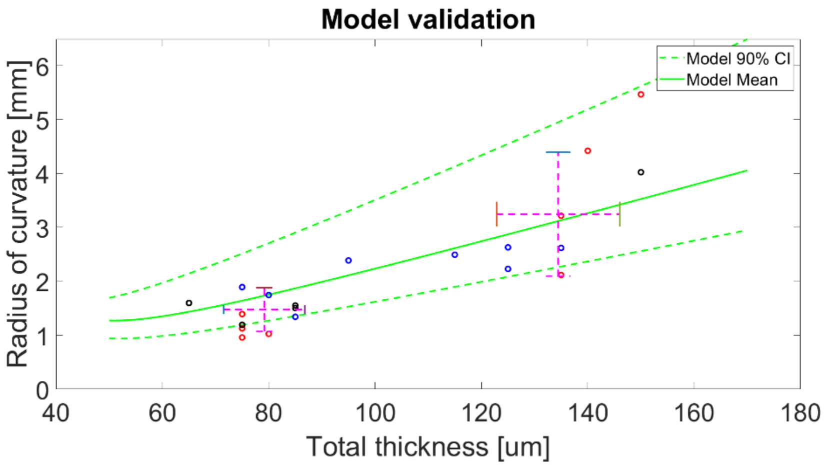


Figure SI3: Third validation of the analytical model according to the global dimension of the structure. Green line indicates the predictive values from the analytical model, whereas the dashed green lines indicate the model 90% confidence interval (CI). Dots represent the experimental measurements. The magenta dashed lines represent the std of the experimental measures, along the radius and along the thickness.

*SI3 Scaffold Scalability*

The capability to fabricate samples with different radii of curvature was evaluated to show the scalability of the developed self-folding scaffold. Exploiting the analytical model, the geometrical parameters needed to obtain a radius of curvature similar to trachea radius of: i) mouse and guinea pig model (~0.75 mm [S6-S7]); ii) human newborn and rabbit model (~ 2.5 mm [SI8-SI9-SI10]); iii) 12-14 years old teenager (~5.5 mm [SI8]); iv) human adult and young pig (~ 7.5-10 mm [SI11-SI12-SI13]) were obtained.

Then, bilayer films were fabricated according to 2.2 and actuated as in 2.3.

Figure SI4 showed the ability to the developed 4D bioprinted scaffold with different radius of curvature (from the bigger to the smaller approximately equal to 7.2, 5, 3, 1.6, 1.1, 0.7 mm) able to match different tracheal sizes of animal models and human ages.

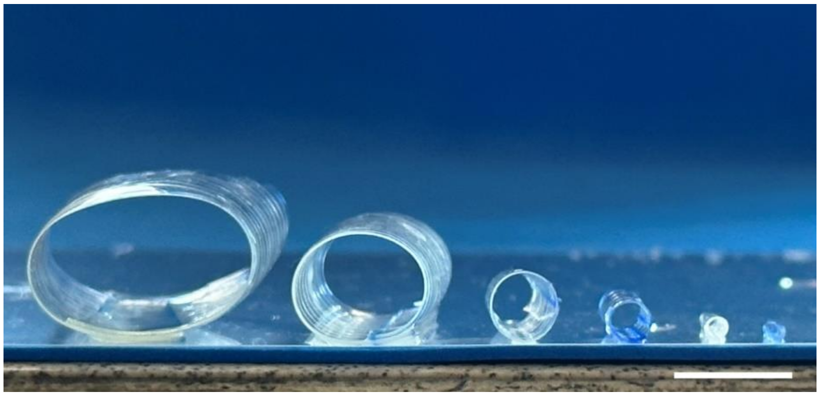

Figure SI4: Scalability of the developed 4D bioprinted self-folding scaffold. Scale bar: 10 mm.

*S4. Holders Fabrication and Testing*

As shown by the actuation tests, as soon as the film is dipped into water it quickly self-folds in few minutes. Since this is not compatible with cell attachment, which on average requires 12 hours, customized holders with open central cavity (Figure SI5A-B) were designed and fabricated to delay the film actuation and to allow cell attachment to the film in its initial flat position. Practically speaking, the holder is placed on the scaffold in a multiwell plate, then cells are seeded while the holder keeps the scaffold flat. After cell attachment (approximately 24h), the holder is removed and the scaffold with the seeded cells self-folds to its final 3D tubular shape (Figure SI5C). The capacity of the scaffold to maintain its shape-shifting behavior after being kept flat by the holder was tested up to 48h. In their final version, the holders were fabricated via indirect AM in polydimethylsiloxane (PDMS), a biocompatible silicone-based organic polymer, widely used for TE application. Before cell culturing, holders were sterilized via ethanol washing (70% v/v in deionized water) for 2 h and 30 mins, three washes with PBS 1X (10 mins each) and finally dried under the hood overnight.

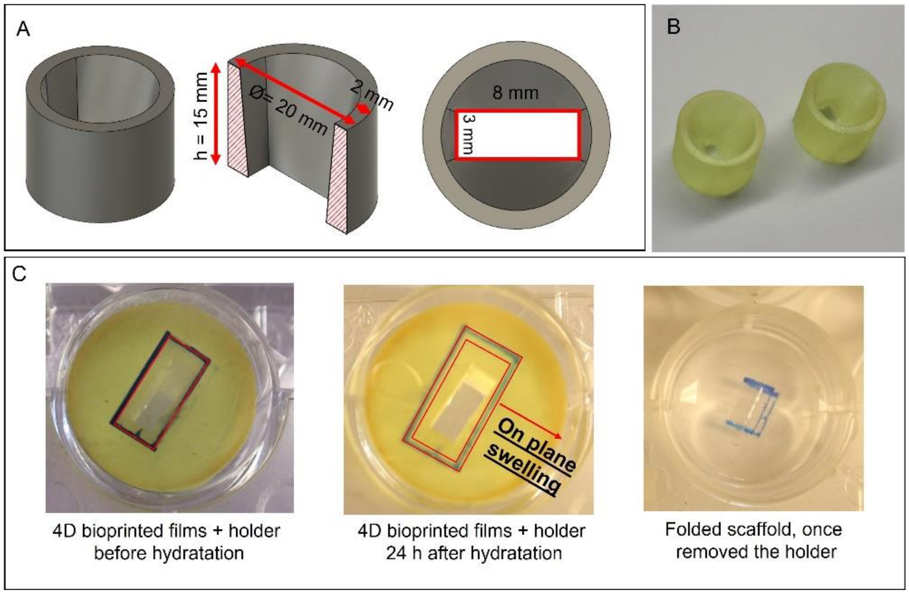


*Figure SI5: Customized holder to delay the scaffold folding. A) CAD model of the holder with main geometrical dimensions. B) Photo of the holders. C) Main steps in the use of the holders*

*SI5: Evaluation of the strain perceived by cells seeded on the self-folded scaffold.*

The strain produced by the scaffold during the folding, and, as a consequence, perceived by attached eCPCs is evaluated and quantified with the Euler-Bernoulli theory. Since the radius of curvature is much greater than the thickness of the structure, the Euler-Bernoulli theory can be applied. Under these conditions, the strain on the external edge of the structure, where eCPCs are seeded, can be obtained by the Equation SI15, derived from Navier' Equation regarding stress experienced by bent beams.

$$\boldsymbol{\varepsilon} = \frac{\boldsymbol{h_2}}{\boldsymbol{\rho}} \quad \text{Eq. SI15}$$

Where $h_2$ is as expressed in Equation SI12.

Due to the non-linearity of the model, the strain produced on the external face of the scaffold depend on the geometry used. Taking as an example a scaffold with a total thickness of 90μm (before the actuation with a theorical $h_2 \approx 60$μm) and a radius of curvature of 1.5 mm (Table 4) the strain evaluation results in an $\varepsilon \cong 6.47\%$.

*SI6. Video of the self-folding of the samples in PBS*

*SI7: Video of the force analysis (figure 7)*

*SI7.1: 20mg*

*SI7.2: 70 mg*

*SI/.3 80 mg*

*SI8: References*

[SI1] Lapomarda, A., *et al.* "Pectin as rheology modifier of a gelatin-based biomaterial ink." *Materials* 14.11 (2021): 3109.

[SI2] Gunasekaran, H., *et al.* "Fabrication and characterization of gelatin/carbon black–based scaffolds for neural tissue engineering applications." *Materials Performance and Characterization* 8.1 (2019): 301-315.

[SI3] Fortunato, G. M., *et al.* "Electrospun structures made of a hydrolyzed keratin-based biomaterial for development of in vitro tissue models." *Frontiers in bioengineering and biotechnology* 7 (2019): 174.

[SI4] Carrabba, M., *et al*., (2016). Design, fabrication and perivascular implantation of bioactive scaffolds engineered with human adventitial progenitor cells for stimulation of arteriogenesis in peripheral ischemia. Biofabrication, 8(1), 015020.

[SI5] Fortunato, G. M., *et al.* "An ink-jet printed electrical stimulation platform for muscle tissue regeneration." Bioprinting 11 (2018): e00035.

[SI6] Kishimoto, Keishi, and Mitsuru Morimoto. "Mammalian tracheal development and reconstruction: from in vivo and in vitro studies." Development 148.13 (2021): dev198192.

[SI7] Amiri, Mohammed H., and Giorgio Gabella. "Structure of the guinea-pig trachea at rest and in contraction." Anatomy and embryology 178 (1988): 389-397.

[SI8] Griscom, N. Thorne, and M. E. Wohl. "Dimensions of the growing trachea related to age and gender."*American Journal of Roentgenology* 146.2 (1986): 233-237.

[SI9] Loewen, M. S., and D. L. Walner. "Dimensions of rabbit subglottis and trachea." Laboratory animals 35.3 (2001): 253-256.

[SI10] Han, Mi-Na, Joong-Hyun Kim, and Seok Hwa Choi. "Evaluation of biomechanical properties and morphometric structures of the trachea in pigs and rabbits." *In Vivo* 36.4 (2022): 1718-1725

[SI11] Judge, Eoin P., et al. "Anatomy and bronchoscopy of the porcine lung. A model for translational respiratory medicine." American journal of respiratory cell and molecular biology 51.3 (2014): 334-343.

[SI12] Breatnach, Eamann, Gypsy C. Abbott, and Robert G. Fraser. "Dimensions of the normal human trachea." American Journal of Roentgenology 142.5 (1984): 903-906.

[SI13] Kamel, Kirollos Salah, Gabriel Lau, and Mark D. Stringer. "In vivo and in vitro morphometry of the human trachea." Clinical Anatomy: The Official Journal of the American Association of Clinical Anatomists and the British Association of Clinical Anatomists 22.5 (2009): 571-579.
